# Isotope Labeling Reveals Complex Microbial Interactions during *Agaricus bisporus* Compost Colonization

**DOI:** 10.64898/2026.08.25.747027

**Authors:** Mădălina M Vîță, Femke van Dam, Michiel VM Kienhuis, Desmond D Eefting, Klaas GJ Nierop, S Emilia Hannula, Lubos Polerecky, Francien Peterse, Jack J Middelburg

## Abstract

Microbial interactions strongly influence carbon and nitrogen flows in mushroom compost, yet their functional roles during *Agaricus bisporus* colonization remain unresolved. We combined PLFA-SIP and nanoSIMS imaging with ITS amplicon sequencing to follow resource flows and microbial activity across spatial scales. Stable-isotope tracers (¹³C- glucose and ¹⁵N-ammonium) revealed that *A. bisporus* simultaneously facilitates and suppresses bacterial populations: fungal activity increased glucose assimilation by bacteria yet reduced overall bacterial biomass. NanoSIMS visualized nutrient-rich microenvironments along hyphae where bacterial ¹³C and ¹⁵N assimilation was elevated. Sequencing showed the fungal community to comprise essentially two organisms, *A. bisporus* and *Mycothermus thermophilus*, which differ approximately elevenfold in their content of the fungal biomarker C18:2ω6,9c. Total fungal PLFA therefore tracks which of the two dominates as much as it tracks fungal biomass. Together these findings reveal coupled fungal–bacterial nutrient processing and show that biomarker-based estimates of fungal biomass require community composition to be known. Multi-scale isotope probing provides a framework for resolving microbial interactions in complex detrital systems.

**Key Points:**

- *A. bisporus* raises bacterial glucose uptake but lowers bacterial biomass
- Bacteria near hyphae show elevated ^13^C and ^15^N uptake at single-cell scale
- Fungal PLFA reflects the combined signal of fungus identity and biomass

**Graphic abstract:** 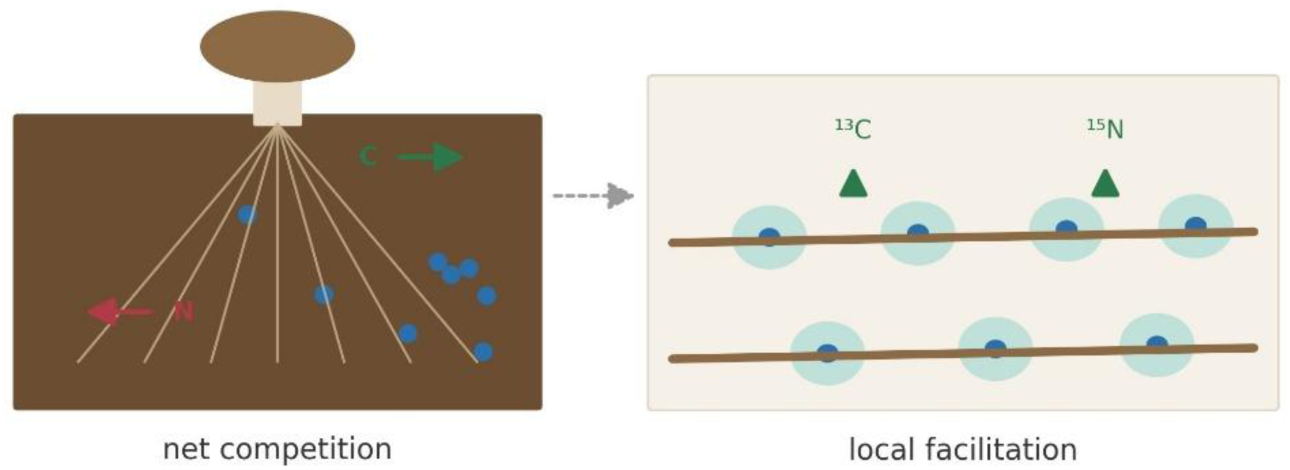

## Introduction

Global mushroom and truffle production exceeded 40 million tonnes in 2021 (Zhao et al. 2025), with the button mushroom *Agaricus bisporus* accounting for nearly half of the market share. The industrial cultivation of *A. bisporus* on a waste-derived substrates also has a high circularity potential (Grimm and Wösten 2018). Therefore, optimising yield and cultivation practices is of significant interest.

The waste-derived substrate is a compost obtained from the fermentation of wheat straw, manure and gypsum which undergoes a multistep composting process prior to its usage. The compost microbiome plays a crucial role from substrate preparation (phases 0–II) through colonization (phases III–IV) to fruiting (Straatsma et al. 1994; Kertesz and Thai 2018). Over time, composting methods have been refined to create optimal growth conditions (Braat et al. 2022). In phase III (PIII), the compost microbial community interacts with the mycelium, influencing growth, yield, and substrate utilisation. For instance, beneficial bacteria such as *Bacillus*, *Streptomyces*, and *Pseudomonas* spp. can enhance mushroom yield and quality by increasing nutrient availability, suppressing pathogens, and promoting mycelial growth and fruiting (Ahlawat and Manikandan 2015; Carrasco et al. 2020; Aydoğdu et al. 2021). Conversely, some moulds can negatively affect growth (Houbraken et al. 2018; Kosanović et al. 2021) .

Bacteria from compost can serve as a nutrient source for *A. bisporus*, which can grow on living or dead bacterial biomass alone (Fermor and Wood 1981; Fermor et al. 1991). Vos et al. (Vos et al. 2017) found that *A. bisporus* presence reduced bacterial biomass. Moreover, lignocellulosic degradation, e.g., of wheat straw, is a multistep process requiring enzymes from multiple microorganisms (Durrant et al. 1991; van der Wal et al. 2013). While *A. bisporus* produces many exoenzymes for straw degradation, bacteria such as *Streptomyces* spp. and *Cellulomonas* spp. also contribute by producing cellulases and hemicellulases (Godden et al. 1989; Yoon et al. 2008).

Although compost microbes clearly influence *A. bisporus* growth, their net contribution to nutrient acquisition during PIII remains unquantified. Most studies have examined microbial community composition during composting and mushroom growth, or the effects of individual strains on yield. These have identified dominant taxa and potential beneficial or harmful mechanisms (Braat et al. 2022), but the cumulative impact on nutrient availability to *A. bisporus* during PIII is poorly understood.

Nutrient flows, particularly of carbon and nitrogen, have been studied in various environments using stable isotope labelling and biomarker analysis (Boschker and Middelburg 2002; De Kluijver et al. 2014; Hannula et al. 2017; Vîta et al. 2026). Phospholipid-derived fatty acids (PLFAs) are useful biomarkers for identifying organisms and estimating microbial biomass (Boschker and Middelburg 2002; Frostegård et al. 2011; De Kluijver et al. 2014; Vos et al. 2017). Combined with stable isotope techniques, PLFA analysis can elucidate food web structure and function (Boschker and Middelburg 2002; Middelburg 2014; Bai et al. 2016). However, PLFA analysis has limitations in compost studies. Taxonomic resolution is restricted to distinguishing fungi, Gram-positive, and Gram-negative bacteria. It can trace carbon and hydrogen but not nitrogen, and it provides only bulk biomass estimates without revealing small-scale spatial variation, such as differential activity within mycelial networks or spatial co-occurrence of bacteria and fungi. Additionally, the fungal biomarker C18:2ω6,9c is present in PII-end compost before *A. bisporus* inoculation, though it declines over time (Vos et al. 2017). Some of these limitations can be addressed with nanoscale secondary ion mass spectrometry (nanoSIMS), which provides spatially resolved isotope data at single-cell resolution, including carbon and nitrogen isotopes (Nuñez et al. 2018). In addition, stable isotope probing of DNA (SIP-DNA) offers higher taxonomic specificity (Friedrich 2006; Dunford and Neufeld 2010; Hannula et al. 2017). When organisms assimilate isotope-labelled substrates (e.g., ¹³C-glucose), the tracer is incorporated into DNA, which can then be density-separated and sequenced to identify tracer-assimilating taxa.

This study examined interactions between *Agaricus bisporus*, compost, and compost-associated microbes during phase III and the onset of phase IV using ¹³C-glucose and ¹⁵N-ammonium chloride as tracers. Substrate utilisation was assessed via changes in total organic carbon and nitrogen and through CO₂ emissions, while carbon transfer to bacteria and fungi was traced using PLFA-SIP. Density separation of DNA was attempted to identify tracer-assimilating taxa but did not sufficiently resolve labelled from unlabelled fractions (Supplementary Results); ITS2 amplicon sequencing of these fractions is therefore used to follow the composition of the fungal community, and nanoSIMS resolved cell-level carbon and nitrogen isotope distributions. Because the fungal PLFA biomarker C18:2ω6,9c is not specific to *A. bisporus*, we also tested whether it reliably reflects *A. bisporus* biomass in a community containing a second dominant fungus.

## Methods

### Incubation experiments

#### Axenic incubations

Axenic cultures of *Agaricus bisporus* and *Mycothermus thermophilus* (CBS 622.91) were grown to characterize and quantify fungal PLFAs. Each culture was incubated separately in Petri dishes at 25 °C and 80% relative humidity on sterile compost agar prepared from homogenised, freeze-dried phase II-end compost as described by Vos et al. (Vos et al. 2017). Sterile polypropylene filters (5 cm diameter, Merck) were placed on the agar to support fungal growth, and incubations continued for three to four weeks until the fungi had fully colonised the filter surface.

#### Phase III-compost incubation

Incubation experiments with *A. bisporus* strain A15 (Lambert Spawn©) were conducted at Utrecht University over 24 days, covering phase III and extending into phase IV of commercial production (Fig. S1), without the addition of a casing layer (Fig. S2A). Mesocosms were prepared with 20 g wet weight phase II-end compost (CNC Grondstoffen©) under four conditions designed to vary initial spawn density and microbial biomass. In the first treatment (1S), five spawn grains were added (1.5% wet weight; industry standard). The second treatment (0.5S) contained two spawn grains supplemented with three sterile rye grains (0.6% wet weight). The third treatment (0S) received no spawn. In the fourth treatment (1S-A), compost was autoclaved three times at 121 °C for 30 min before inoculation with five spawn grains.

At the start of incubation, all treatments received 1.1 mg ¹³C-glucose (U-¹³C₆, ¹³C atom fraction 99%; Cambridge Isotope Laboratories) and 0.05 mg ¹⁵N-ammonium chloride (¹⁵N atom fraction 99%) per gram of compost, suspended in 500 µl sterile deionised water and applied as small droplets. Each 20 g mesocosm therefore received 22.0 mg glucose (0.118 mmol), corresponding to 0.709 mmol ¹³C, and 1.0 mg ¹⁵N-ammonium chloride, corresponding to 0.019 mmol ¹⁵N. To assess whether simple carbon sources were limiting, two additional triplicate treatments (1S* and 0.5S*) were prepared in parallel, following the same inoculation as 1S and 0.5S but with the isotopic mixture reapplied after 14 days, together with 11.4 mg unlabelled glucose per gram of compost. Control samples with unlabelled glucose and ammonium chloride were included for each treatment to establish baseline isotopic values and ensure equal nutrient content across treatments.

All incubations were carried out in the dark at 22 °C and 80% relative humidity. Destructive sampling took place after 3, 7, 14, and 24 days, with an additional sampling at day 17 for the 1S*, 0.5S*, 1S, and 0.5S treatments (Fig. S3A). Carbon dioxide production was measured over a 1 h period using a GC-0006-W ExplorIR® CO₂ sensor and GasLab® software, followed by trapping in sodium hydroxide solution for isotopic analysis using a GasBench (Thermo Scientific™). Measurements were performed in triplicate. Cultures were then freeze-dried, homogenised using a Herzog HP MA/HP PA mill, and analysed for bulk carbon and nitrogen concentrations and isotopic composition using an elemental analyser coupled to an isotope ratio mass spectrometer (EA-IRMS; Thermo Fisher Scientific). The same homogenised material was used for PLFA extraction (Supplementary Fig. S3A).

#### Phase III and IV-focused square-dish incubation

In parallel, isotope-labelling experiments (dedicated to SIP-DNA) were performed in square Petri dishes (10 × 10 cm; Greiner Bio-One B.V.) containing 45 g phase II-end compost (CNC, Milsbeek, The Netherlands) inoculated with ten spawn grains of *A. bisporus* strain A15 (Sylvan, Horst, the Netherlands) (1.5% wet weight). At the start of incubation, 0.74 mg glucose was added per gram of compost; half the dishes received ^13^C-labelled glucose (99%), while the remainder received unlabelled glucose as controls (Fig. S2B). Each 45 g dish therefore received 33.5 mg glucose (0.18 mmol), corresponding to 1.08 mmol ¹³C in the labelled dishes. No ¹⁵N tracer was applied to this incubation.

Colonization of the compost represented phase III, and samples collected during this period are referred to as phase III compost. After full colonization at day 14, 20 g peat-based casing (CNC) was added. The spawn run from day 0 to 21 was conducted at 22 °C and 80% relative humidity, after which mushroom formation was induced by venting at 18 °C and 80% relative humidity until day 30. Duplicate compost and casing samples (5 g each) were collected at days 0, 7, 14, 21, and 30 for PLFA and DNA analyses, with DNA stored at −20 °C (Fig. S3B).

### PLFA extraction and analysis

PLFAs were extracted using a modified Bligh and Dyer method (Bligh and Dyer 1959), as described by de Kluijver et al. (de Kluijver et al. 2021), and analysed by gas chromatography–combustion–isotope ratio mass spectrometry (GC-c-IRMS; Delta Plus XP, Thermo Finnigan). Detailed extraction and analytical procedures are provided in the Supplementary Methods. The PLFAs used as biomarkers are listed in Table S1. Isotope ratios and PLFA concentrations were processed in R using the Rlims package. Incorporation of ^13^C into PLFAs was calculated following Boschker and Middelburg (2002) and expressed as atom fraction x(^13^C), representing the proportion of tracer incorporated relative to total carbon in the biomarker. In tracer addition experiments, growth was inferred from increases in x(^13^C) due to labelled substrate incorporation and decreases due to growth on unlabelled background resources, assuming constant total biomass (Fig. 1).

**Figure 1.**
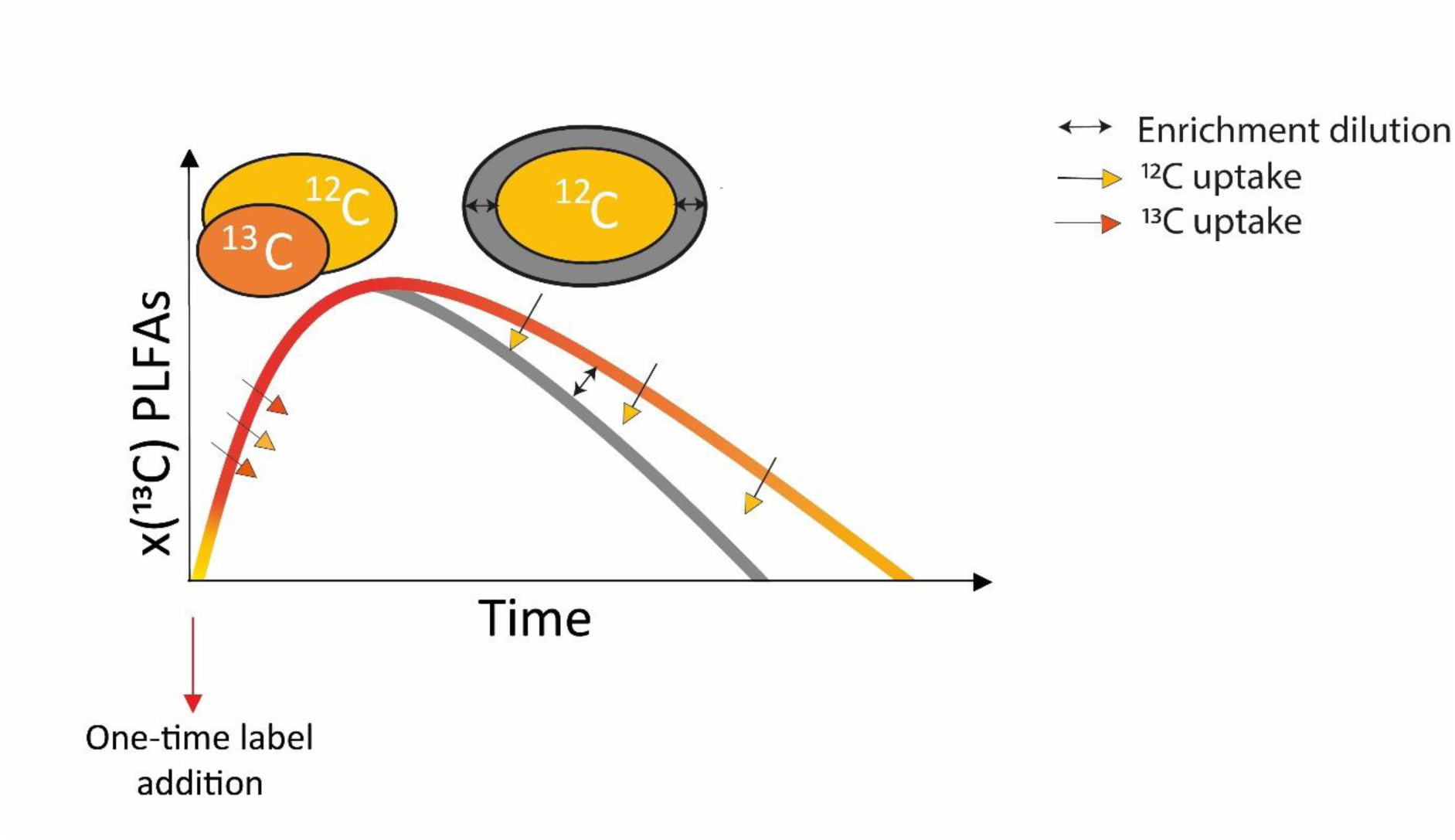
Conceptual model illustrating tracer (^13^C) incorporation into PLFAs over time. Immediately after tracer addition, rapid uptake increases PLFA ^13^C atom fraction x(^13^C). As biomass turns over, incorporation of unlabelled (^12^C) substrate decreases relative enrichment. The grey line illustrates a scenario of higher carbon-use efficiency, where more ^12^C is incorporated per unit turnover.

### DNA extraction and ITS2 sequencing

DNA was extracted, fractionated on CsCl density gradients, screened by quantitative PCR, and the ITS2 region amplified and sequenced on an Illumina MiSeq (paired-end 250 bp); full procedures are given in the Supplementary Methods. The density separation did not sufficiently resolve labelled from unlabelled DNA, principally because fungal labelling was too low for a detectable density shift (Supplementary Results), so the sequencing data are reported as community composition across fractions.

### Nanoscale secondary ion mass spectrometry (nanoSIMS)

NanoSIMS was used to resolve microscale patterns of carbon and nitrogen assimilation by fungal hyphae and associated bacterial cells during compost colonization. NanoSIMS imaging enabled quantification of ^13^C and ^15^N incorporation at the single-cell level, allowing discrimination between fungal hyphae, attached bacteria, and free-living bacterial cells. Imaging targeted compost incubations receiving ^13^C-glucose and ^15^N-ammonium chloride to assess spatial heterogeneity in isotope incorporation within fungal–bacterial interaction zones. Detailed sample preparation, imaging conditions, region-of-interest selection, and isotope ratio calculations are described in the Supplementary Methods.

### Statistical analysis

Statistical analyses of PLFA concentrations, CO₂ production, and bulk carbon and nitrogen measurements were performed in R v4.2.0. Differences among treatments and sampling time points were evaluated using one-way analysis of variance (ANOVA) followed by Bonferroni post hoc tests when overall effects were significant. Student’s t-tests were used for pairwise comparisons where appropriate. Prior to analysis, data were tested for normality and homogeneity of variances. When ANOVA assumptions were violated, Kruskal–Wallis rank-sum tests were applied, followed by Dunn’s post hoc comparisons with Benjamini–Hochberg correction for multiple testing. For datasets in which variances were not homogeneous, as assessed by Bartlett’s test, chi-square and Kruskal–Wallis-derived p-values were reported instead of F-values and ANOVA p-values. All statistical tests were two-sided, and significance was assessed at α = 0.05. Full statistical outputs, including test statistics and adjusted p-values for all comparisons, are provided in Supplementary Tables S2–S4.

## Results

### Compost physical-chemical changes and tracer dynamics

The total nitrogen, carbon, and water contents remained relatively stable during the incubation period regardless of the conditions (Fig. S5), with none of the variations being statistically significant (Table S2 and S3). Among them, the autoclaved condition showed very high variability in results for carbon and water content (Fig. S5 B and C).

Quantification of the bulk tracer in the compost according to the different treatments revealed variability in time and treatment for both ^13^C and ^15^N. The initial ^13^C content of the compost was variable (Fig. 2A). However, this variability was not statistically significant (Table S2, [F(2,3)=2.88, p=0.167]). Moreover, the values were close to the 0.709 mmol of ^13^C added to the compost (Fig. S6). On the contrary, from the 0.019 mmol ^15^N added to each culture, only a third was detected at day 0, and the ^15^N content was significantly lower than that measured at any later time point (Fig. 2B). Freeze drying may have caused the loss of ^15^N-ammonia during sample processing, and the conversion of ^15^N- ammonium into ^15^N-biomass during incubation consequently led to nitrogen conservation. The amount of ^13^C from the added glucose decreased over time as part of it was respired as ^13^CO_2_ (Fig. 2A and S7A).

**Figure 2.**
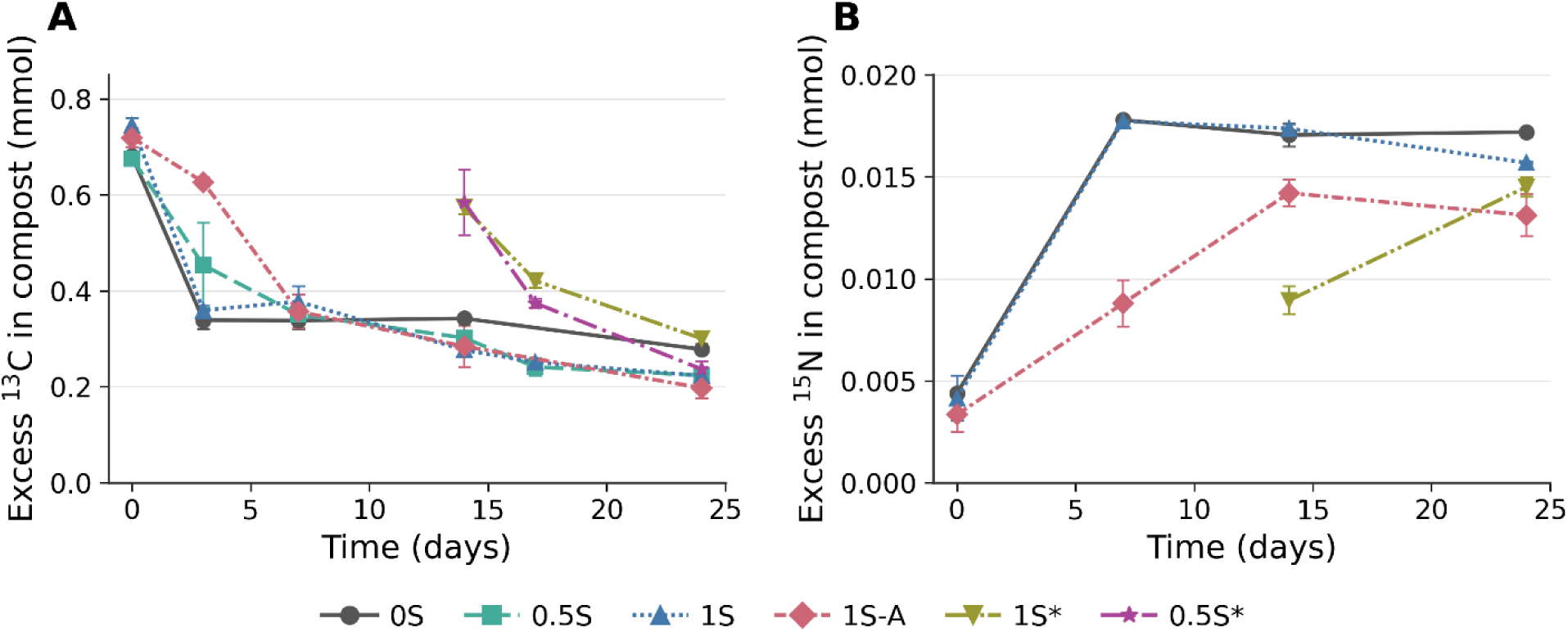
^13^C and ^15^N excess of the compost**. ( A)** excess of ^13^C over time (days of incubation). (**B)** excess of ^15^N in bulk compost over time (days of incubation). Error bars represent minimum and maximum values (n = 2).

The autoclaved 1S-A compost initially showed the slowest ^13^C loss from the compost, as also reflected in limited ^13^CO_2_ release (Fig. S7A), consistent with the quantity of tracer initially remaining in this compost. On day 3, there was more ^13^C in 1S-A than in any other condition, although from day 7 on, no significant differences were detected (Table S2). Therefore, autoclaving caused a major initial reduction in microbial activity. Moreover, the autoclaved treatment had significantly lower ^15^N than the other treatments considered after 24 days of incubation (Table S3, [F(2,3) =9.127, p=0.029] (Fig. 2B), but lower amounts of ^15^N were also detected at day 0. The mesocosm without *A. bisporus* (0S) eventually contained more ^13^C than the other mesocosm. However, this difference was not statistically significant. In 1S* and 0.5S* the tracer was applied on day 14 rather than at the start, together with 11.2 mg of unlabelled glucose per mesocosm to test whether the community was substrate-limited. This lowered the ¹³C enrichment of the glucose pool approximately 1.5-fold, while the ¹⁵N addition was undiluted. Despite this, and despite having ten days rather than twenty-four in which to assimilate the tracer, by day 24 these two conditions contained amounts of ¹³C and ¹⁵N comparable to the conditions labelled at day 0 (Fig. 2), and interpolation of the cumulative curves showed that they had the highest ¹³CO₂ production by day 24 (Fig. S7). Glucose added to compost in which a fungal network was already established was therefore metabolised more rapidly than glucose added at inoculation.

To summarise, although the various treatments contained variable amounts of tracer, from the joint measurements of ^13^C in the compost and ^13^CO_2_ production, we can conclude that the introduction of *A. bisporus* into the compost caused a more rapid consumption of the compost’s glucose over time than did the endogenous microbial community alone.

### Biomass composition

The Fungal:Bacterial PLFA ratio in the treatment without *A. bisporus* remained stable at approximately 0.1 throughout the incubation, showing no major shift in the microbial community composition (Fig. 3A). The highest F:B ratio was calculated under autoclaving conditions. Overall, in the other treatments, the presence of *A. bisporus* affected the F:B ratio only after two weeks, when the fungi, presumably *A. bisporus*, became more dominant, suggesting therefore an inhibitory effect, i.e., growth inhibition or predation, of the mushroom on the bacterial population. For mesocosms that started with low spawn density (0.5S), the increase in fungal PLFAs relative to bacterial PLFAs started later than that in high spawn density ones (1S), i.e., at 17 days rather than at 14 days.

**Figure 3.**
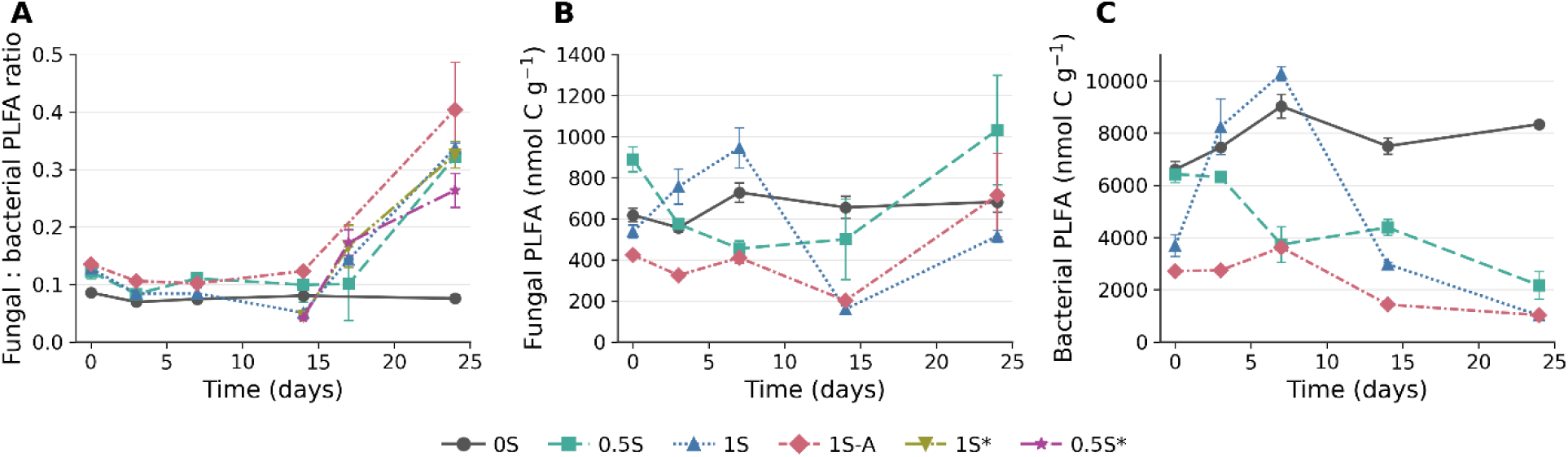
PLFA-derived biomass indicators across treatments. (**A)** Fungal:Bacterial PLFA ratio. (**B)** Fungal PLFA concentrations. (**C**) Bacterial PLFA concentrations. The ratio was calculated from the PLFA concentrations of the biomarkers listed in Table S1. The fungal biomarker was divided by the sum of all bacterial PLFAs. Error bars represent minimum and maximum values (n = 2).

The initial fungal and bacterial PLFA concentrations differed among 0S, 0.5S, and 1S at day 0 [F (2,3) =17.58, p=0.022] (Fig. 3B and C, Table S4 and S5), complicating biomass comparisons among treatments at different time points. PLFA concentrations were also variable at T0 in the other treatments (Fig. S8 A and B). Although both groups of biomarkers were variable, their ratios were rather constant during the first 0-14 days (Fig. 3A), indicating that fungal and bacterial PLFAs in compost covaried.

In the square-dish incubation, both total bacterial PLFAs (3954 ± 321 nmol C g^-1^) and fungal PLFA (382 ± 54 nmol C g^-1^) were present in high quantities at the start of the experiment (Fig. S9A) followed by an increase over the initial 7 days, and then a decrease after day 14. After this time point, similarly to 1S bottle incubation (Fig. 3A), bacterial PLFAs declined further, approaching approximately 30% at the end of the incubation, whereas fungal PLFA increased by a factor of approximately 4. In the casing samples, the total bacterial PLFAs concentration was lower than that in the compost (Fig. S9C). The biomarker levels of both groups increased over time. Between days 14 and 31, the total bacterial PLFAs concentration increased by approximately 2-fold to a final concentration of 214 ± 92 nmol C g^-1^, whereas the fungal PLFA started from undetectable concentrations and reached 452 ± 292 nmol C g^-1^ at day 30.

### 13C dynamics in biomass

The fraction of ^13^C to total carbon content of PLFAs shifted in time for both bacteria and fungi in all treatments (Fig. 4A-F). Under the conditions in which the label was added on day 0 of the incubation, namely 1S, 0S, and 1S-A (Fig. 4 A, B, and C), the bacterial PLFAs had a minimum 10-fold and maximum 93-fold higher atom fraction compared to the fungal ones. Therefore, the results indicate a more actively growing and dividing bacterial population. The highest atom fraction value was recorded for the autoclaved compost, where the microbial community was particularly low at the beginning of the incubation (Fig. 4C), probably because of reduced competition for resources and more tracer availability per microorganism. Moreover, 1S-A exhibited a lag phase in label incorporation in both the fungal and bacterial populations.

**Figure 4.**
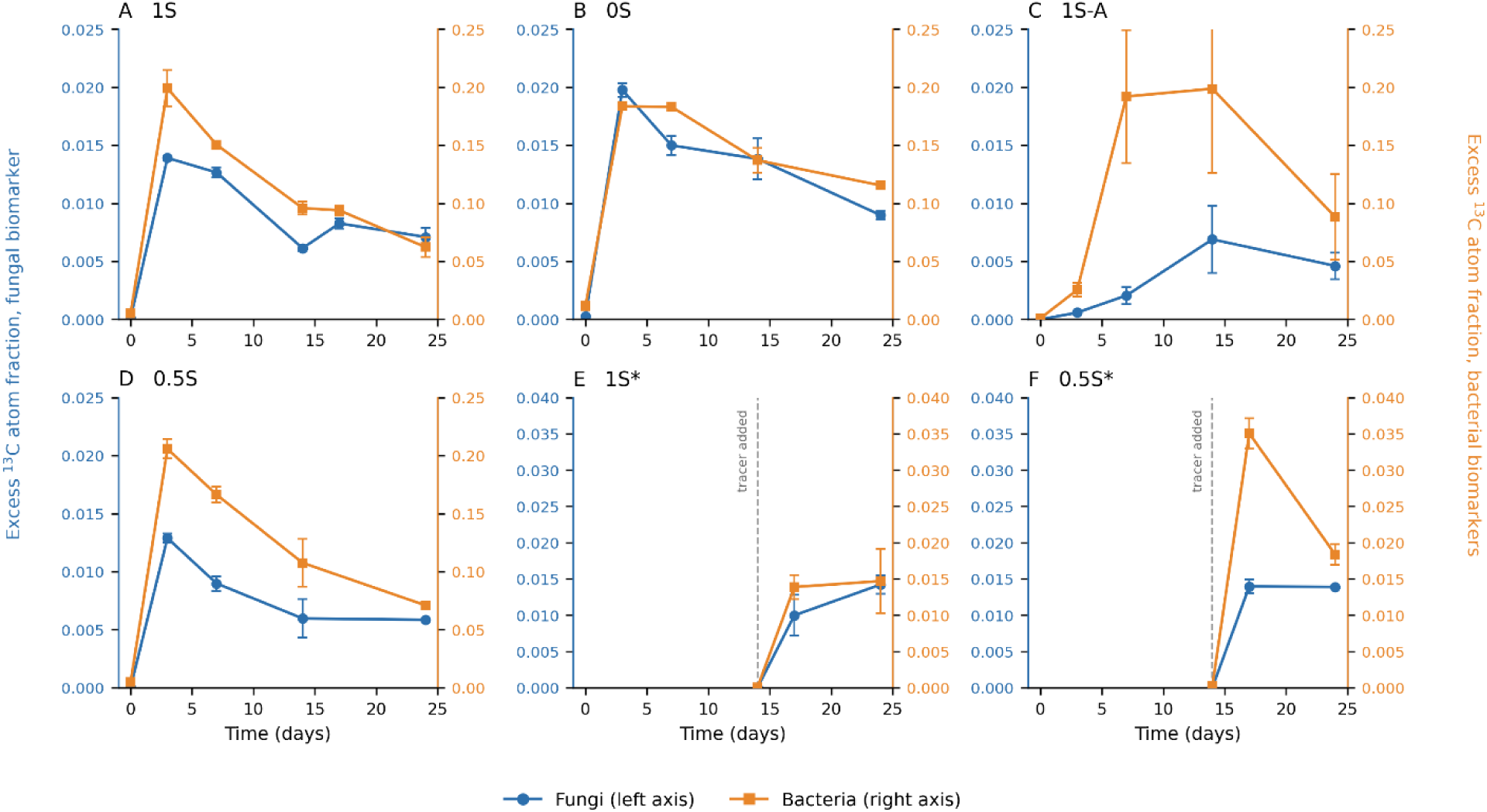
Excess ^13^C atom fraction of fungal and bacterial PLFA biomarkers over time in each treatment: (A) 1S, (B) 0S, (C) 1S-A, (D) 0.5S, (E) 1S*, (F) 0.5S*. Blue circles: fungal biomarker, left axis; orange squares: bacterial biomarkers, right axis. Note the different axis scaling in panels E and F, where the dashed vertical line marks tracer addition on day 14. Labels denote the initial set-up of the cultures (1S = standard spawn density; 0.5S = reduced density; 0S = no spawn; * tracer added day 14; 1S-A = autoclaved substrate). Error bars represent minimum and maximum values (n = 2).

The trends in atom fraction over time also differed for each treatment. The decrease in the atom fraction of bacteria under standard conditions (1S) was steeper than that in the treatment without *A. bisporus* (0S). Because the label was introduced into the system as a one-time addition, a more rapid loss indicates the incorporation of non-labelled substrates (Fig. 1). In 1S*, where the tracer was added after 14 days, the bacterial community was less actively growing with respect to the treatment in which label was added on day 0 (1S). The atom fraction of the bacteria had a similar low fraction as the one of the fungal group. However, the decrease in activity was less marked under the 0.5S* condition (Fig. 4 E and F). This shows that at 14 days, the bacterial community size is not only diminishing as seen in the previous section (Fig. 3B), but it is also less actively growing, with this last trend being spawn density dependent. Moreover, the highest fungal ^13^C atom fraction was recorded in 0.5S*, where in addition to the reduced competition due to loss of activity in bacteria, there are also fewer fungal hyphae; therefore, more tracer is available per unit of biomass.

In the square-dish incubation, the excess fraction of ^13^C of compost bacteria was much higher than that of fungi, indicating a more active community in tracer uptake, i.e., faster and more incorporation of carbon per total carbon unit. Moreover, the excess ^13^C atom fraction of bacterial PLFAs peaked on day 7 and then decreased, whereas that of the fungal PLFA continuously increased until the last sampling day (Figure S9B). In contrast to observations in the compost, the bacterial and fungal PLFAs in the casing had similar excess ^13^C atom fractions at 21 days, indicating similar activities, and diverged near the end when the bacterial excess ^13^C atom fraction further increased while that of the fungi did not (Fig. S9D).

### Zoom on fungal community dynamics

Analysis of fungal communities in compost and casing was carried out for the square-dish incubation. DNA was fractionated on CsCl density gradients before sequencing. At the labelling achieved here, 0.52 atom% excess ¹³C in the fungal biomarker (Fig. S9B, D), the expected shift in DNA buoyant density is about 0.0002 g/mL, well below the resolution of the gradient. Fractions are therefore treated as subsamples of each gradient and the sequencing data are reported as community composition rather than as evidence of ¹³C assimilation (Supplementary Results).

Across all sequenced fractions the community was dominated by two genera, *Agaricus* and *Mycothermus thermophilus*, which together accounted for 99.8% of fungal reads. A single genus made up more than 90% of the reads in 54 of the 64 fractions recovered inside the gradient. Variability among replicates was high from day 0 and persisted through the early stages of incubation. Substrate was the only significant structuring factor in a PERMANOVA on Bray-Curtis distances (R2 = 0.070, F = 4.79, *p* = 0.034; Fig. S17).

In the casing, *M. thermophilus* was scarce at 21 days, with *Agaricus* dominating all fractions (Fig. 5A). Excluding *Agaricus* (Fig. S10) revealed a more diverse community, particularly at day 21, when Chao1 and Shannon indices indicated a rich and even structure (Fig. S11A). By day 30 this diversity had declined as *Agaricus* increased. Where *Agaricus* was excluded, *M. thermophilus* was the most abundant taxon in most fractions, exceeding 95% relative abundance.

**Figure 5.**
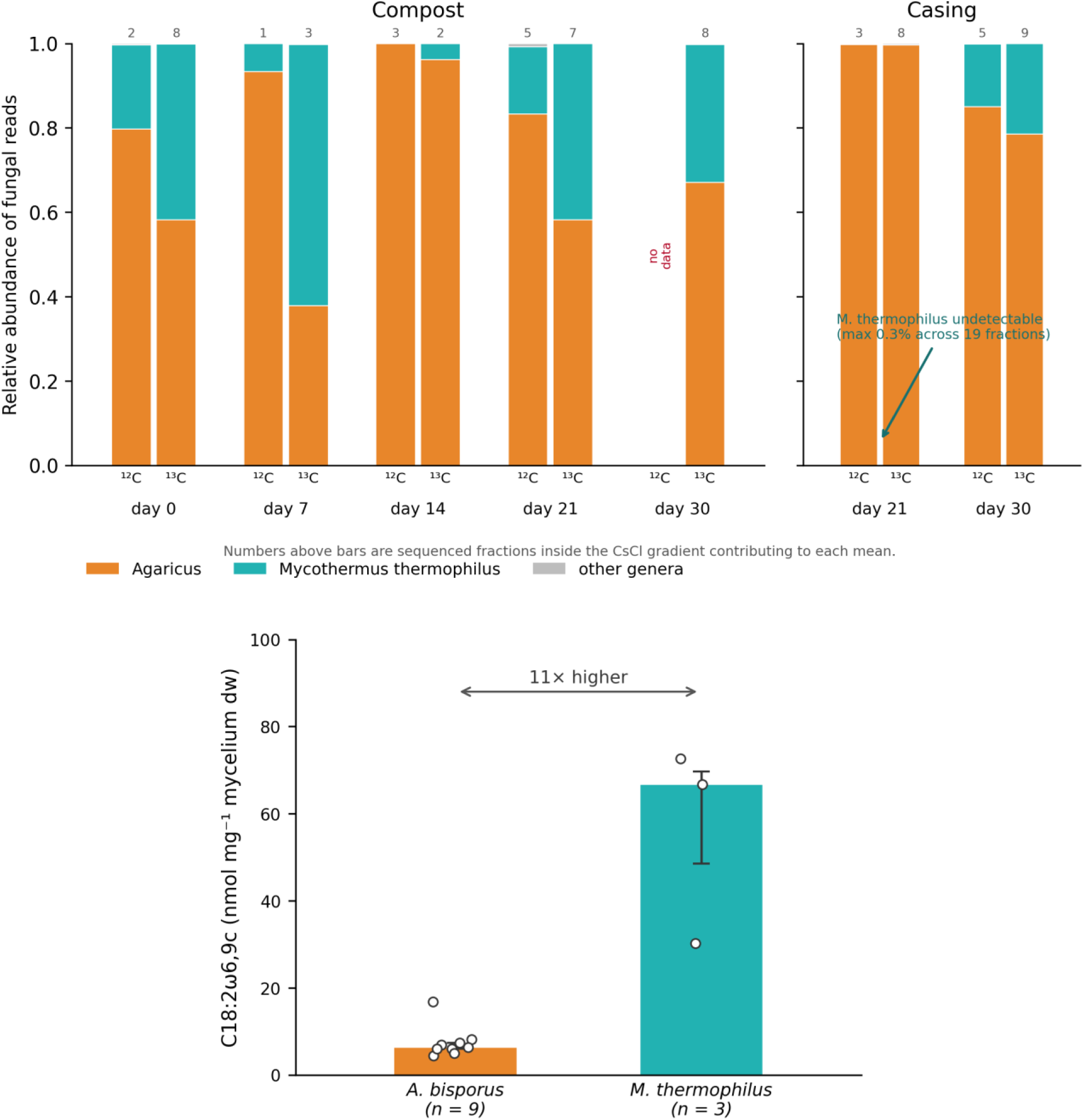
(A) Fungal community composition in compost and casing over the incubation. Bars show the mean relative abundance of fungal reads across sequenced fractions inside the CsCl gradient, for unlabelled (¹²C) and labelled (¹³C) dishes separately; numbers above bars give the number of contributing fractions. Agaricus and M. thermophilus together account for 99.8% of all fungal reads. M. thermophilus was undetectable in the casing at day 21, seven days after the casing was applied, reaching at most 0.3% across 19 fractions, and was present in both unlabelled and labelled dishes by day 30. No unlabelled compost fractions from day 30 fell inside the gradient. (B) Linoleic acid (C18:2ω6,9c) content of the two dominant species in axenic culture. Bars show the median, error bars the interquartile range, and open circles the individual cultures (n = 9 for A. bisporus, n = 3 for M. thermophilus). Medians are reported rather than means because each group contains one culture that departs markedly from the others. M. thermophilus contains approximately 11 times more linoleic acid per unit dry weight than A. bisporus (66.8 against 6.3 nmol mg⁻¹), so total fungal PLFA is weighted towards M. thermophilus and a change in which of the two dominates alters the PLFA signal independently of total fungal biomass.

The clearest contrast is in the casing. Seven days after it was applied, *M. thermophilus* did not exceed 0.3% of fungal reads in any of the 19 casing fractions sequenced; by day 30 it exceeded 70% in four of 18 fractions, in three of the four gradients and in both labelled and unlabelled dishes. With four gradients per timepoint the increase is not statistically resolvable (Mann-Whitney *p* = 0.12), but the appearance from absence is unambiguous. *M. thermophilus* therefore colonised the casing during cropping.

Other genera appeared sporadically, including *Scopulariopsis* in compost at 7 days and *Penicillium*, *Aspergillus* and *Apiotrichum* in casing at 30 days (Fig. 5A). *Apiotrichum* and *Saitozyma* were briefly prominent in the casing at day 21 before *Agaricus* and *M. thermophilus* predominated by day 30. Overall, these results show a community structured by *A. bisporus*, with greater diversity when this genus is excluded, particularly in the casing (Fig. S11A), and competitive dynamics between *A. bisporus* and *M. thermophilus* (Figs. S10 and S11B).

Because the fungal community reduces to these two organisms, interpretation of the fungal PLFA biomarker depends on which of them dominates. Linoleic acid (C18:2ω6,9c) is used as a fungal biomarker on the assumption that it is consistently present in fungi and absent from other organisms and organic pools. Neither assumption holds fully here: linoleic acid was detected in the fruiting body, in the rye grains used for spawn production and in the spawn itself, with very similar chromatograms for all three (Fig. S12). Cells of *M. thermophilus* contained a median of approximately 11 times more linoleic acid than those of *A. bisporus*, 66.8 against 6.3 nmol mg⁻¹ dry weight of mycelium (Fig. 5B). Both species showed a strong positive correlation between C18:2ω6,9c content and mycelial dry weight (Fig. S13); fitting through the origin, since a culture of zero biomass contains no PLFA, gives 6.7 nmol mg⁻¹ for *A. bisporus* against 62.1 nmol mg⁻¹ for *M. thermophilus*. Linoleic acid content is therefore relatively constant within a strain but varies considerably between species, so the living biomass of *M. thermophilus* has a disproportionate influence on total fungal PLFA. These comparisons are approximate: the two species were cultured over non-overlapping biomass ranges, 7.0 to 41.5 mg dry weight for *A. bisporus* against 4.1 to 6.2 mg for *M. thermophilus*, and content per unit dry weight rises with culture size in both, so *M. thermophilus* is if anything underestimated relative to *A. bisporus*. Full statistical output for all PLFA comparisons is given in Tables S2 to S4.

### Nanoscale isotopic imaging

Imaging of microorganisms on polycarbonate filters revealed that no hyphae were detected after 7 days of incubation under any of the four conditions (Fig. 6, 7 days). However, multiple fields of view (FOVs) did contain bacterial cells. Hyphae were found only after 14 days of incubation under conditions containing *A. bisporus* (Fig. 6, 14 and 24 days).

**Figure 6.**
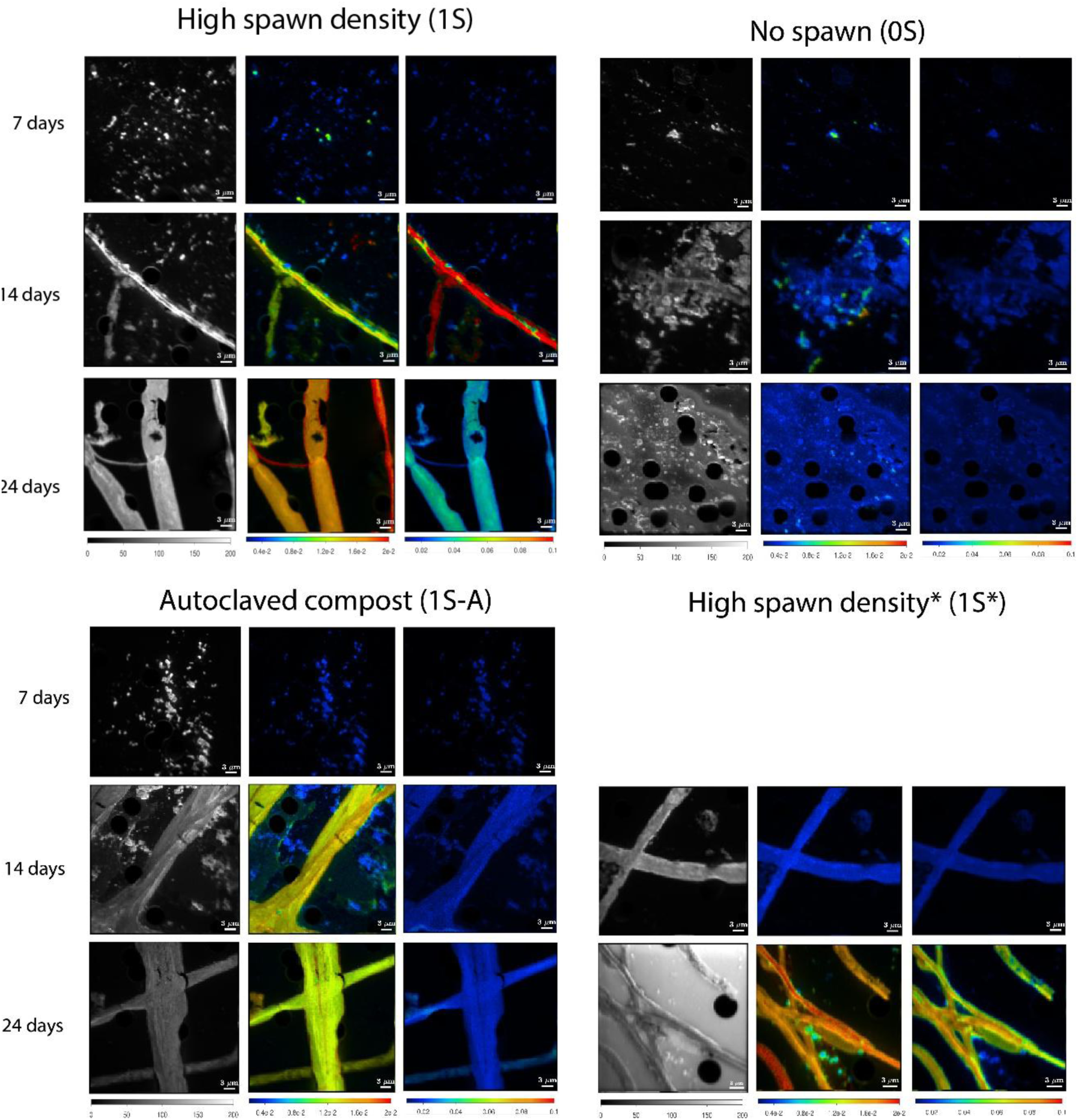
Fields of view from nanoSIMS imaging. Each group of panels represents an incubation condition, and the titles indicate the type of compost under different conditions. Each row shows a different ratio: left: carbon image, ^12^C_2_/plane/pixel; middle: nitrogen enrichment, ^12^C^15^N/(^12^C^14^N+^12^C^15^N); right: carbon enrichment, 0.5*^12^C^13^C/(^12^C2+0.5*^12^C^13^C). The rows represent time points, from top to bottom: 7, 14, and 24 days. Colour intensity, dark blue to red, indicates the intensity of the signal and mass abundance, with dark blue representing the minimum and red the maximum of the chosen scale.

Overall, the ^15^N atom fractions increased over time in both bacterial populations (attached and free-living) and *A. bisporus* hyphae (Fig. 7B). In contrast, bacteria and *A. bisporus* were less enriched in ^13^C at 24 days than at 14 days in conditions labelled on day 0, such as 1S, 1S-A, and 0S (Fig. 7A). This decline in ^13^C enrichment may be attributed to the progressive loss of ^13^C label as ^13^CO_2_ and the incorporation of ^12^C-rich organic matter (Fig. 1, 4).

**Figure 7.**
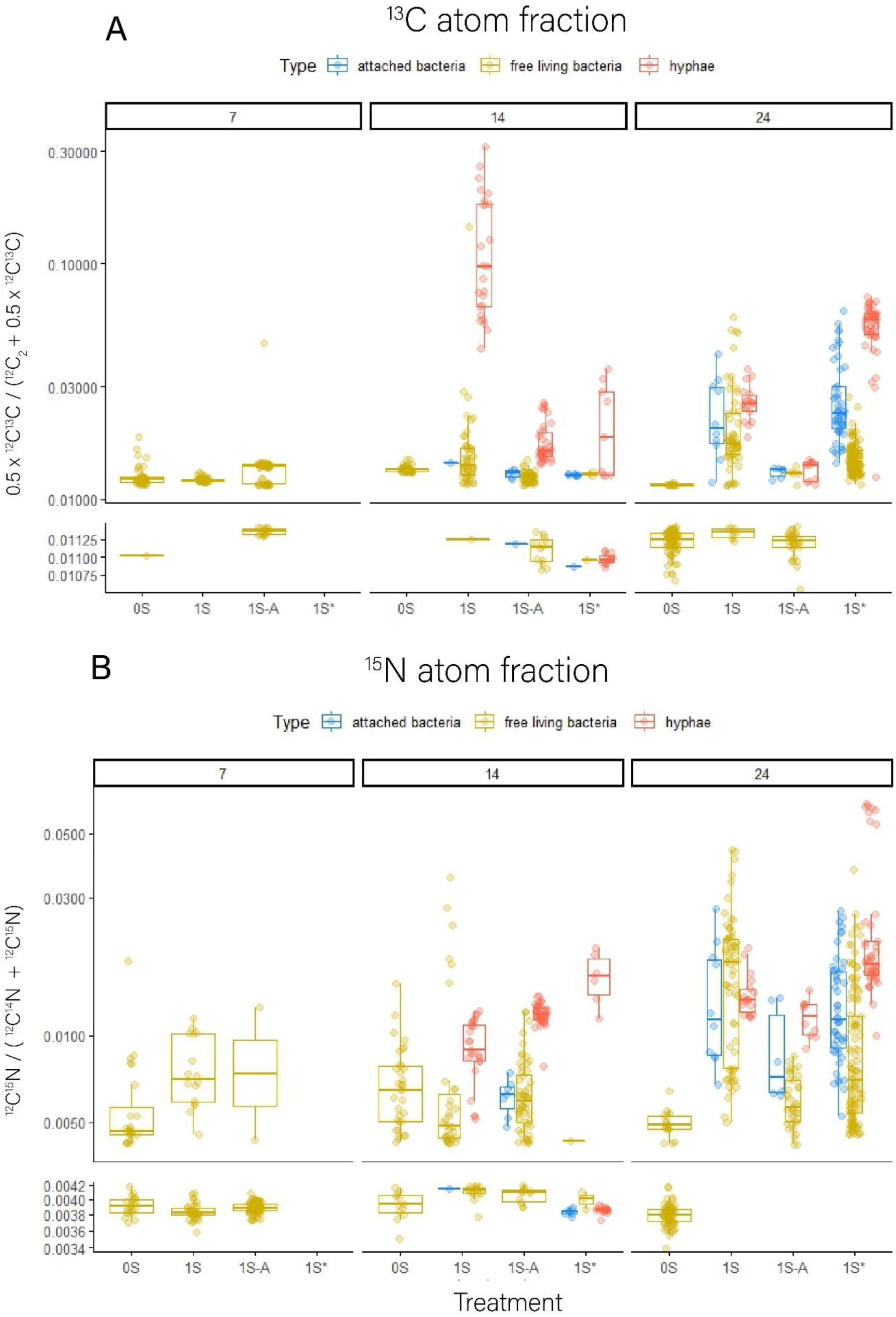
nanoSIMS imaging: regions of interest (ROI) among the different treatments. The number above each group indicates the sampling time point (7, 14 and 24 days). The different colours differentiate the microorganisms imaged (attached bacteria, free living bacteria, and hyphae). **A** ^13^C and **B ^15^**N atom fraction. The upper panel presents the measurements of labelled cell, whereas the bottom panel presents the measurements of non-labelled cells. The range for the unlabelled cells was established based on the atom fraction of the cells in the non-labelled treatment. The Y- axis shows the atom fraction of the ROIs and is displayed in log scale. The relative abundances of each class of cells per treatment and time are presented in Figure S12.

In autoclaved compost, the hyphae were less enriched in ^13^C than in the non-autoclaved condition at 14 days (Fig. 6, Autoclaved and Fig. 7A) (K-W, p=4.80E-17). Conversely, ^15^N enrichment was higher in autoclaved conditions at 14 days than in 1S (K-W, p=7.19E-10), likely due to reduced competition for resources. By the end of the incubation period, ^15^N enrichment in the autoclaved treatment was on average, lower than that in the non-autoclaved condition, although this difference was not statistically significant.

Image analysis showed that not all bacterial cells were co-enriched in ^13^C and ^15^N (Fig. S14). Some samples contained only one tracer despite homogeneous mixing before labelling. Nonetheless, a moderate to strong correlation was observed for bacterial cells in 1S at 7 and 24 days, 1S* at 24 days, and 1S-A and 0S at 14 days (R²=0.48-0.87). Given the small sample size and the restricted area, it is possible that bacterial carbon and nitrogen uptake are generally correlated. Additionally, not all bacterial cells were enriched (Fig. S15). For the enriched bacteria, the average ^13^C and ^15^N atom fractions at 14 and 24 days were lower in the 0S treatment than in the conditions containing *A. bisporus* [K-W, p=1.54E-35 and ANOVA, p=6.07E-07]. In the presence of *A. bisporus*, bacterial atom fractions were closer to hyphal values (Fig. 7 A and B, attached vs. free-living), suggesting that hyphae may create a nutrient-rich microenvironment. In support of this finding, more attached bacterial cells were detected in 1S and 1S* on the surface of the hyphae after 24 days compared with 14 days (Fig. S15). Correlation analyses of ^13^C and ^15^N for *A. bisporus* hyphae revealed no correlation (Fig. S14), indicating that the two tracers were utilised differently by the hyphae.

A loose attachment was observed between fungal filaments and bacteria when imaging in higher resolution 400 depth planes (Fig. S16). Bacterial cells were primarily found on the outer layer of the hyphae (Fig. S16, OL). This layer exhibited a looser and flatter structure compared to the core of the filament, together with irregular edges and a lack of the strong, defined edges seen in the hyphal core. This type of attachment could allow bacterial cells to move along the hyphal length partially explaining why there was migration of ^13^C enriched bacteria from the compost to the casing in the square-dish incubation (Fig. S9D).

## Discussion

### Inhibition and competition in compost habitats

Given the initial variability in PLFA content among different treatments, a comparison of the biomarker concentrations was not possible. However, some consistent trends were observed within different time points for the same treatment. Specifically, in the treatments that contained mushrooms, the bacterial PLFA concentrations decreased over time, whereas in the treatment without the spawns, the PLFA concentrations remained constant (Fig. 3B). This agrees with the trends observed by Vos et al. in which bacterial PLFAs decreased in time when *A. bisporus* was present (Vos et al. 2017).

In addition to this loss of bacterial biomass (Fig. 3B , Fig. S9), treatments in which the tracer was added after 14 days of incubation revealed a decline in bacterial activity as well. Specifically, the fraction of ^13^C to total carbon of bacterial PLFAs was 10-fold lower than that of their day 0 labelled counterpart when a high spawn density was used and five-fold lower in the case of low spawn density (Fig. 4). This implies that the bacterial community incorporated less label in the presence of *A. bisporus*. As mentioned above, this could be partly due to a decrease in bacterial cell activity. However, the rapid uptake of the ^13^C-glucose by *A. bisporus* also limited the availability of this resource to the bacterial population. Nevertheless, when considering atom fraction in the same analysis, the fungal ^13^C incorporation was higher than expected only in the treatment with low spawn density. This could be explained by the lower hyphal density, resulting in more glucose availability per unit of mushroom biomass. In conclusion, a decrease in bacterial activity in the presence of *A. bisporus* is highly possible. The decrease in bacterial activity and biomass in the presence of *A. bisporus* indicates that fungi suppress other microbiota, but it does not conclusively prove that they predate on bacteria, consume microbes for nutrients, or inhibit the growth of bacteria via competition for resources. Moreover, the different degrees of inhibition among treatments suggest that this inhibitory effect is dependent on spawn density.

Due to the non-specific nature of the fungal PLFA biomarker, no significant correlations between fungal and *A. bisporus*-specific trends were discerned. The process of scaling up PLFA contents to total biomass is complex, as the PLFA content in cell membranes varies among organisms, including fungi (Fig. S13; see also Camenzind et al. 2024(Camenzind et al. 2024)) and bacteria (Boschker and Middelburg 2002). The average linoleic acid content in *M. thermophilus* was up to 11 times higher than that in *A. bisporus*. It is known that the linoleic acid content in certain fungi varies with life stage and temperature (Haubert et al. 2008). Furthermore, Rousk et al. reported no significant alterations in C18:2ω6,9c content during incubations at varying pH levels, despite a fivefold increase in fungal biomass(Rousk et al. 2010). They proposed that some variability in PLFAs could be ascribed to the adaptation of cell membrane composition in response to pH changes, as documented in previous studies (Šajbidor 1997; Rousk et al. 2010). This underscores the intricate and dynamic nature of lipid composition in fungi, which is influenced by a range of environmental and biological factors.

While the use of PFLA as biomarker resolves bacterial from fungal dynamics, it is restricted to ^13^C and does not provide cell-specific and spatial information. Consequently, nanoSIMS was an essential analytical tool for addressing this gap. Under conditions where labelling was conducted later in the incubation (1S*), the hyphae of *A. bisporus* exhibited the highest levels of ^15^N compared to any other condition (Fig. 7). Although not the highest, the ^13^C atom fraction under the same conditions (1S*) was also notably high. This indicates that *A. bisporus* efficiently assimilates nutrients when fully developed, consistent with the high respiration rate of the ^13^C-glucose under the same conditions (Fig. S7). However, the bacterial cells in this treatment were not particularly low in ^13^C and ^15^N. When distinguishing between the attached and free-living bacterial populations, higher ^13^C and ^15^N enrichments were observed in the attached group, suggesting that the hyphae create a nutrient-rich microenvironment around them. Given that only bacteria in proximity to the hyphae were imaged, the nanoSIMS results did not directly contrast with the PLFAs-based F:B ratio, which suggested a less active bacterial population. Although the same analysis did not show a particularly high fungal enrichment with nanoSIMS, we confirmed that *A. bisporus* specific activity was indeed higher than that observed in the initial fungal population of the compost. In addition to the fungal-bacterial competition, image analysis revealed that some bacterial cells were not co-enriched in ^13^C and ^15^N (Fig. S14), despite the tracers being homogeneously mixed. This could be a result of competition and differences in affinity for tracers among the bacterial cells forming the community.

### Bacterial-fungal cooperation

Indications of cooperation between the bacterial and fungal populations were visible at various levels, from community properties (e.g., CO_2_ production) to microorganism-specific activity. Higher levels of ^13^CO_2_ were released from the treatments labelled on day 14, although the label was in place for a shorter period (Fig. S7). Moreover, the lowest ^13^CO_2_ production rate occurred in treatments in which no *A. bisporus* was present or the initial microbial community was reduced by autoclaving. Therefore, glucose was respired faster when both *A. bisporus* and the microbial community of the compost were present.

The isotope dilution of labelled PLFA biomass provided additional evidence that non-labelled, background organic substrates were assimilated at higher quantities (Fig. 4). Excess ^13^C, i.e. the ratio of labelled over total bacteria PLFAs, showed that bacteria in presence of *A. bisporus* lost label faster than those in the absence of the mushroom. This indirectly indicates that more organic carbon was available for uptake by bacteria when *A. bisporus* was present. Consistently, nanoSIMS imaging showed greater enrichment of both ^13^C and ^15^N in bacteria at the end of the incubation when hyphae were present. As proposed above, this suggests the presence of a more active microenvironment around the hyphae, specifically increased nutrient availability due to the presence of *A. bisporus*, which stimulates the activity and growth of mycobiomes (Frąc et al. 2022). In support of this finding, more tracer was present in the bacteria in the treatments where *A. bisporus* was present in the nanoSIMS image results. However, this was not observed in the atom fractions of bacterial PLFAs in treatments with no *A. bisporus*. Therefore, more fields of view need to be imaged to support this possibility.

Strong evidence emerged on how the bacterial community or parts of it benefits from the presence of *A. bisporus*. However, this relationship is not one-sided, as some results indicated that *A. bisporus* also benefits from a microbial population in compost. The hyphae in the autoclaved treatment were less enriched in ^13^C than in the non-autoclaved counterparts at all time points. Given the reduced initial fungal and bacterial activity in 1S-A (Fig. 4C), higher amounts of tracer were expected. A higher hyphal density could explain this, as less tracer would be available per unit of biomass, although extremely high fungal PLFA concentrations were not measured in this treatment that could support this possibility. Although nanoSIMS imaging showed that fungal hyphae were less enriched in ^13^C in the autoclaved treatment, the excess ^13^C ratios based on nanoSIMS imaging were similar to the fraction of labelled PLFAs compared to total PLFAs (∼ 0.01). We can, therefore, assume that the biomarker used to quantify fungi (i.e., linoleic acid: 18:2ω6,9) was produced by fewer organisms in the autoclaved compost than in the other treatments.

The combined PLFA-DNA fungal biomarker approach, while limited in providing precise quantitative data, offered insights into the dynamics of the fungal community. The total DNA pool was divided into multiple density fractions, but not all were sequenced, leaving some DNA unaccounted for and making it challenging to determine whether the observed changes in relative abundances were due to a reduction in *M. thermophilus* or an increase in *A. bisporus* (Morton et al. 2019). Given the active growth of *A. bisporus* during the cropping process, as observed visually, changes in relative abundance were likely primarily driven by growth. However, the low fungal PLFA content on day 14 (Fig. S9A) could indicate either that *A. bisporus*, with its lower PLFA content, became dominant or that there was a significant reduction in *M. thermophilus* biomass. Based on only the PLFA data, it is not possible to distinguish between these two alternatives. The ITS data resolve this. At matched buoyant density, *M. thermophilus* accounted for 39% of fungal reads in compost at day 0, fell to 3% at day 7 and 2% at day 14, and recovered to 52% by day 21. Because *M. thermophilus* carries approximately 11 times more linoleic acid per unit dry weight than *A. bisporus*, a compositional swing of this magnitude lowers total fungal PLFA without any loss of fungal biomass. Hence, the day 14 fungal PLFA minimum is at least partly due to a change in fungal community composition and does not reflect only a decline in fungal biomass. This illustrates the limitations in using PLFA data to quantitatively reconstruct fungal dynamics.

A shift in the fungal population of the casing samples was detected over time, first dominated by *Agaricus bisporus* and then by *M. thermophilus*. The enrichment of the fungal biomass in the casing compared with the compost suggests active migration and growth of both fungi in the casing. This is further supported by the results of PLFA analysis, which indicated that fungal biomass was more concentrated in the casing than in the compost, implying that fungi migrated into the casing. Previous studies have underscored the importance of *M. thermophilus* in the optimal growth of *A. bisporus,* particularly during substrate preparation (López et al. 2021). The present work adds evidence that this beneficial cooperation between the two species is not restricted to phase II (compost preparation) but extends into phase III (vegetative growth of *A. bisporus*).

In addition to *Agaricus* and *Mycothermus,* several other genera were found in the compost. These genera were detected sporadically and were outliers to typical *M. thermophilus* and *A. bisporus* pairs. Notably, *Scopulariopsis,* which was previously identified in self-heating composts and indoor systems (Samson and von Klopotek 1972; Woudenberg et al. 2017), was present. Specifically, *Scopulariopsis fimicola* is known to cause white plaster mold, which can grow on the compost surface or in casing layers, inhibiting the growth of *A. bisporus* mycelium (Mishra 2011). Additionally, *Aspergillus* and *Penicillium*, ubiquitous in many ecosystems, were also abundantly reported *in A. bisporus* compost (McGee et al. 2017; Kertesz and Thai 2018). Similarly, *Kernia* has been identified in *A. bisporus* compost [46]. At 21 days, the dominant genera in the casing, excluding *A. bisporus*, were *Apiotrichum*, *Saitozyma*, and *Aspergillus*. *Apiotrichum* is frequently found in casing soil and is considered a member of the endogenous fungal community (Taparia et al. 2021; Tello Martín et al. 2022). Although there are no specific reports of *Saitozyma* in the casing or compost used for *A. bisporus* cultivation, it might be related to the *Candida* genus, as *Saitozyma podzolica* was previously classified as *Candida podzolica* (Liu et al. 2015). Overall, the fungal community identified here aligns with current knowledge of the fungal communities in *A. bisporus* compost and casing.

PLFA analysis provided insights into bacterial biomass dynamics, revealing a decrease in bacterial PLFAs in the compost and a simultaneous increase in the casing during incubation. However, different types of bacteria have varying proportions of specific biomarkers in their cell membranes (Ratledge and Wilkinson 1988), and extrapolation from PLFA content to total community biomass introduces uncertainty. Additionally, certain PLFAs, such as i14:0, a/i15:0, cy17:0, and cy19:0, are known to vary in concentration with pH in arable and forest soils (Högberg et al. 2007; Rousk et al. 2010). This is relevant because the pH of *A. bisporus* compost undergoes acidification during the cropping process, dropping from 8 in phase I to 6 by day 16 of phase III (Jurak et al. 2014). Despite these variations, individual bacterial PLFAs showed biomass trends similar to the overall sum of bacterial biomarkers in soil (Jurak et al. 2014). Overall, there are complex interactions between bacterial communities, PLFA biomarkers, and environmental factors like pH that shape biomass dynamics in compost and casing.

The presence of *A. bisporus* may have stimulated endogenous bacteria in the casing, potentially through exudate release. ^13^C enrichment observed in bacteria could be attributed to bacterial feeding on these exudates, consistent with the spatial association between fungal hyphae and bacteria observed in nanoSIMS images of the bottle incubations, which contained no casing layer (Fig. 7). Alternatively, some bacteria migrate from the compost to the casing, potentially using fungal hyphae as a pathway, a mechanism supported by previous studies (Barto et al. 2012). This migration could result in the low ^13^C enrichment observed in the casing’s bacterial population, as the enriched bacteria would be measured alongside the unlabelled bacterial community already present in the casing. NanoSIMS imaging revealed that these bacteria are loosely attached to the outer layer of fungal hyphae, which is primarily composed of β-glucans (Gow 2025). This supports the idea that bacteria can use fungal hyphae as a “highway”, a well-documented phenomenon (Kohlmeier et al. 2005; Warmink et al. 2011; Simon et al. 2015). The ^13^C atom fraction analysis in this study suggests that both mechanisms may be at play: while bacteria and fungi share similar atom fractions by day 21, the bacterial population’s atom fraction increases further from day 21 to day 30. Given that no label was present in the casing, the source of ^13^C enrichment could be exudates from *A. bisporus* or other organisms.

## Conclusions

Isotope labelling combined with PLFA and nanoSIMS is a robust approach for investigating the growth and interactions of bacteria and fungi in compost. Bacterial activity declined after 14 days at high spawn density, while bacterial glucose uptake increased in the presence of *A. bisporus*. NanoSIMS showed increased bacterial activity around hyphae, reduced carbon assimilation by *A. bisporus* in sterilised compost, and distinct nitrogen and carbon uptake patterns between fungi and bacteria, with nitrogen accumulating over time while glucose-derived carbon was rapidly used and dissipated. The study also exposed a limitation of using PLFA as fungal biomarker: C18:2ω6,9c is carried unequally by the two dominant fungi and is present in the spawn substrate, so total fungal PLFA reflects which fungus dominates as much as how much fungal biomass is present. Amplicon sequencing resolved that ambiguity, showing *M. thermophilus* falling from 39% of fungal reads in compost at day 0 to 2% at day 14 and recovering to 52% by day 21, and colonising the casing between days 21 and 30 from undetectable to locally dominant. This shifting dominance rather than displacement points to a dual role of competition and cooperation between *M. thermophilus* and *A. bisporus*.

## Data availability

Raw ITS2 amplicon sequences from the density-gradient fractions are deposited at NCBI under BioProject PRJNA1503199, with associated BioSample and SRA records; these data are held until publication. Processed data, comprising fraction metadata with measured buoyant densities, the ASV table and taxonomic assignments, and PLFA- SIP and nanoSIMS measurements, are archived at Zenodo under DOI 10.5281/zenodo.22094356. The R analysis scripts are archived separately at Zenodo under DOI 10.5281/zenodo.22097242. Both records are cross-linked to BioProject PRJNA1503199. Additional data relating to this work are available in the associated doctoral thesis, DOI 10.33540/2915.

## Supporting information

Supplementary Methods, Figures S1toS17 and Tables S1toS4

