## Supplementary Methods, Figures S1toS17 and Tables S1toS4 for "Isotope Labeling Reveals Complex Microbial Interactions during *Agaricus bisporus* Compost Colonization"

**PLFA extraction and GC-c-IRMS analysis**

Phospholipid fatty acids (PLFAs) were extracted from freeze-dried compost, casing, and axenic fungal material using a modified Bligh and Dyer method (Bligh and Dyer 1959), following de Kluijver et al. (De Kluijver et al. 2014). Approximately 0.3 g of freeze-dried and homogenised material was extracted with 7.5 ml of a 2:1:0.8 (v/v/v) mixture of methanol, dichloromethane, and phosphate buffer. Phase separation was induced by adding additional phosphate buffer and dichloromethane, after which the organic phase was collected.

Total lipid extracts were fractionated on activated silica columns using sequential elution with dichloromethane (apolar lipids), acetone (neutral lipids), and methanol (polar lipids). The polar lipid fraction containing PLFAs was subjected to mild alkaline methanolysis using methanolic NaOH and a toluene:methanol (1:1, v/v) mixture at 37 °C for 30 min in the presence of a C19:0 fatty acid methyl ester internal standard.

The resulting fatty acid methyl esters (FAMEs) were dissolved in hexane, spiked with a C12:0 internal standard, and analysed by gas chromatography–combustion–isotope ratio mass spectrometry (GC-c-IRMS) using a Delta Plus XP isotope ratio mass spectrometer (Thermo Finnigan) coupled to a gas chromatograph equipped with a J&W CP-Sil 5 CB column (25 m length, 0.32 mm internal diameter, 0.12 µm film thickness).

FAMEs were assigned by comparison of retention times with those of an external FAME standard mixture (Supelco 37 Component FAME Mix), supported by equivalent chain length (ECL) values calculated from the retention times of C12:0, C16:0 and C19:0. Identities were confirmed by gas chromatography–mass spectrometry (GC-MS) (Finnigan Trace GC Ultra), and double-bond positions of monounsaturated FAMEs were determined from dimethyl disulfide (DMDS) adducts analysed by GC-MS (de Kluijver et al. 2021). PLFA concentrations were quantified against the C19:0 and C12:0 internal standards, and concentrations and isotope ratios were processed in R using the Rlims package v1.03 (Soetaert et al. 2015). Incorporation of $^{13}C$ into individual PLFAs was calculated following Boschker and Middelburg (2002) (Boschker and Middelburg 2002) and expressed as atom fraction $x(^{13}C)$. The PLFAs used as biomarkers for bacterial and fungal biomass are listed in Table S1 (based on (Matcham et al. 1985; Frostegård and Bååth 1996; Zelles 1997, 1999).

##### DNA extraction, stable isotope probing, and sequencing

DNA was extracted from 0.2 g aliquots of freeze-dried compost and casing samples to characterise the active fungal community. Samples were homogenised using a bead beater (Retsch MM200) and stored at −20 °C prior to extraction using the Quick-DNA Faecal/Soil Microbe Microprep kit (Zymo Research), following the manufacturer's instructions. DNA quality was assessed by electrophoresis on 1% agarose gels, and DNA concentrations were determined using a Qubit HS assay kit (Thermo Fisher Scientific).

Approximately 2 µg DNA per sample was subjected to CsCl density gradient ultracentrifugation following a modified published protocol (Dunford and Neufeld 2010). DNA was dissolved in CsCl gradients and ultracentrifuged for 72 h at 325,913 g and 20 °C. Gradients were fractionated into 18 fractions, which were precipitated using 30% polyethylene glycol in 1.6 M NaCl.

Quantitative PCR targeting the ITS2 region was performed using primers ITS4 (TCCTCCGCTTATTGATATGC) and ITS1F (CTTGGTCATTTAGAGGAAGTAA). This assay was used as a rough presence screen rather than as a quantitative selector. The ITS1F/ITS4 amplicon (approximately 600 to 800 bp) is too long for reliable SYBR-based quantification, quantification was against an *Escherichia coli* standard so no fungal standard curve was available, and no-template controls amplified at cycle thresholds comparable to those of the samples. Cycle threshold values did not distinguish fractions inside the CsCl gradient from those outside it (median 28.6 against 28.3, Mann–Whitney $p = 0.81$; Spearman correlation with buoyant density rho = -0.05, $p = 0.64$), so the screen carried no information about where DNA had banded. A parallel bacterial 16S assay on the same DNA gave 13 to 18 cycles of separation from the no-template controls, indicating that the DNA was suitable for amplification and that the limitation lay with the fungal assay. Fractions were selected for sequencing on the basis of this screen; as set out in the Results, the resulting selection did not give matched density coverage between labelled and unlabelled gradients. PCR amplification was carried out using primers ITS4 and ITS3 (CTAGACTCGTCATCGATGAAGAACGCAG), and amplification quality and ITS2 presence were confirmed by electrophoresis on 1.5% agarose gels. Sequencing was performed at Genome Québec using the Illumina MiSeq platform (paired-end 250 bp).

##### Sequence analysis

Sequence data were processed using the DADA2 ITS pipeline (v1.8) (Callahan et al. 2016), with modifications described by Rolling et al. (Rolling et al. 2022). Primer sequences were removed using cutadapt (Martin 2011). Because ITS length varies among fungal taxa, denoising was performed without fixed-length trimming. Filtering parameters (maxEE = 2, trunQ = 2) were optimised using the FIGARO tool (Weinstein et al. 2019) to maximise read retention.

Taxonomic assignment followed the DADA2 workflow using the UNITE+INSD database with a bootstrap threshold of 50 and seed = 100 (Nilsson et al. 2015). Analyses were conducted in R v4.0.0. Fractions were grouped into heavy (position 1–6), intermediate (position 7–12), and light (position 13–18) classes based on their position in the gradient, not on measured density. Buoyant density (g/mL) was recorded for every fraction; in most gradients the light-position fractions (13–18) fell below 1.690 g/mL, the threshold below which DNA does not band in CsCl, meaning these fractions lay outside the resolvable density window. Because *Agaricus* dominated most samples, a parallel analysis excluding ASVs assigned to this genus was performed.

Alpha diversity was assessed using the Chao1 richness estimator and Shannon diversity index, calculated in QIIME2 (Bolyen et al. 2019) and R.

**NanoSIMS sample preparation and analysis**

Sterilised isopore polycarbonate membrane filters (5.0 µm pore size, 2.5 cm diameter; Merck®) were inserted into one culture bottle per condition and sampling time. Filters from treatments 1S, 1S*, 1S-A, and 0S were collected after 7, 14, and 24 days of incubation (Fig. S2). Discs (0.5 cm diameter) were cut from filters, washed in phosphate-buffered saline, and fixed in 2.5% glutaraldehyde and 4% paraformaldehyde (Electron Microscopy Sciences) at 4 °C for 48 h, followed by post-fixation in 1% osmium tetroxide in $ddH_2O$.

Samples were dehydrated in ethanol at −20 °C (50% and 70% for 1.5 h), at 4 °C (85%, 95%, and 100% for 1 h), and finally at room temperature in 100% ethanol for 1.5 h. Filters were stored in 95% ethanol at −20 °C prior to final dehydration, air-dried, and sputter-coated with gold.

Imaging was performed using a NanoSIMS 50L instrument (Cameca, France) for nominal masses $^{12}C_2$, $^{13}C$, $^{14}N$, $^{15}N$, $^{31}P$, and $^{32}S$. Fields of view were scanned 300–400 times before alignment and accumulation. Image analysis was performed using Look@NanoSIMS (Polerecky et al. 2012). Regions of interest (ROIs) corresponding to fungal hyphae, attached bacteria, free-living bacteria, and exopolymeric substances were selected based on combined phosphorus, carbon, nitrogen, and oxygen signals (Fig. S4). Oxygen-rich particles were identified as oxalic acid crystals and excluded from analysis.

The $^{13}C$ atom fraction was calculated as
$x(^{13}C) = 0.5 \cdot {}^{13}C / ({}^{12}C_2 + 0.5 \cdot {}^{13}C)$,
and the $^{15}N$ atom fraction as
$x(^{15}N) = {}^{15}N / ({}^{14}N + {}^{15}N)$.

Enrichment thresholds were defined as the mean atom fraction of bacterial cells in unlabelled controls plus two standard deviations.

### Supplementary figures

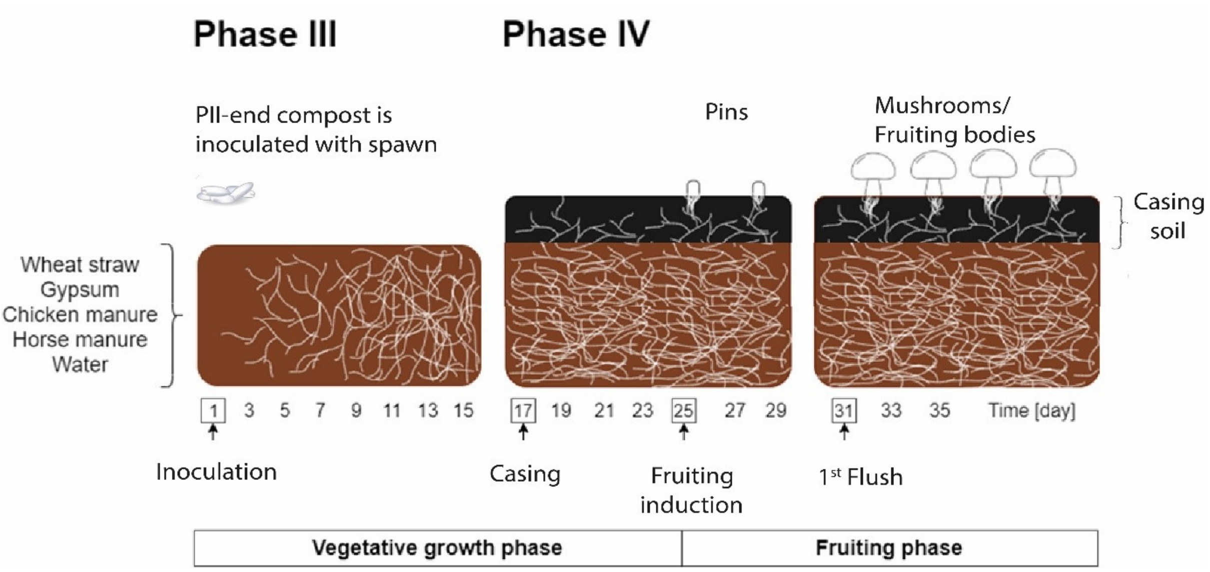


**Figure S1.** Schematic representation of *A. bisporus* cultivation. Spawn was added to the phase II (pII)-end compost in phase III (pIII), and the fungus mycelium colonised the compost. The fruiting phase begins as pins start to form at the surface of the casing layer, which eventually develop into the fruiting bodies (mushrooms).

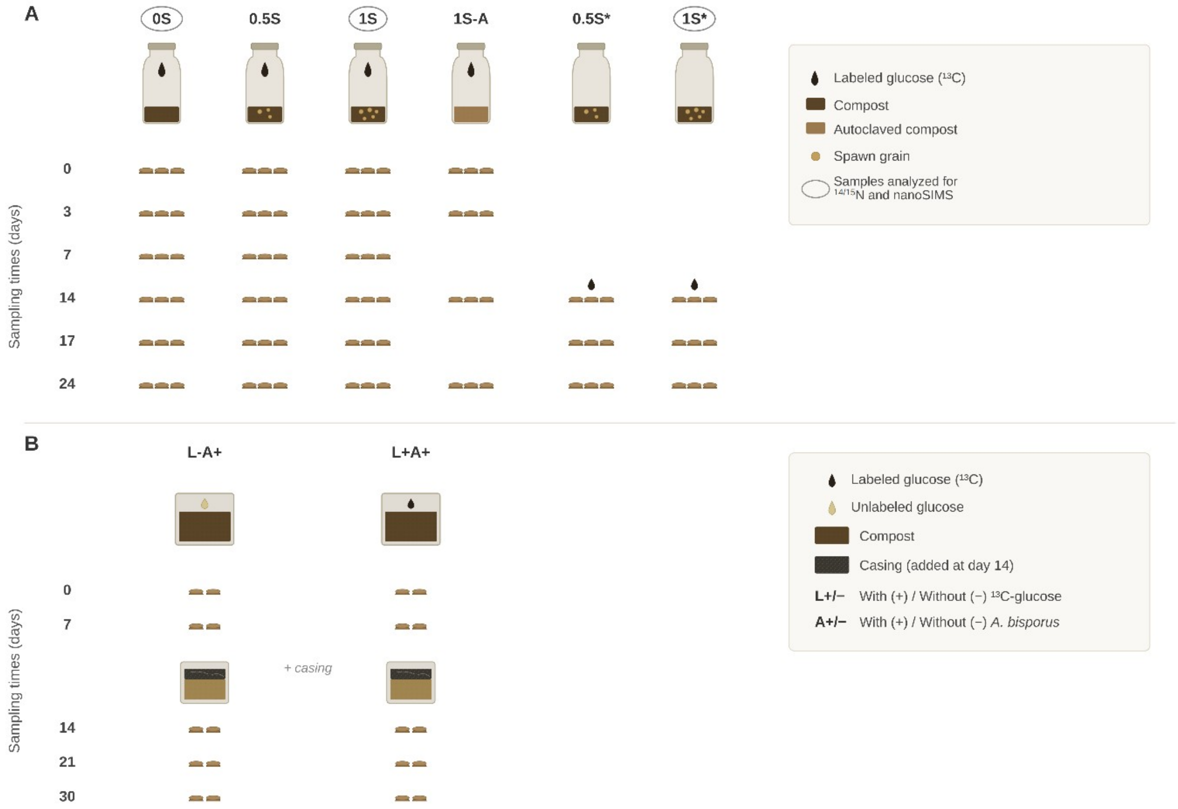

**Figure S2.** Schematic sampling diagram of different conditions and sampling moments in the two different incubations. A: Schematic of compost-only experimental setup. Additional $^{13}C$ background samples on days 0 and 14 are not included in the diagram. B: Schematic of experimental setup of upscaled compost and casing setup. DNA was extracted from duplicate casings and compost separately at different time points (0, 7, 14, 21 and 30 days of incubation). The experiment was conducted under two conditions: L-A+ and L+A+, referring to the absence (-) or presence (+) of a $^{13}C$-glucose label, and A+ indicates the presence of *A. bisporus* in the compost.

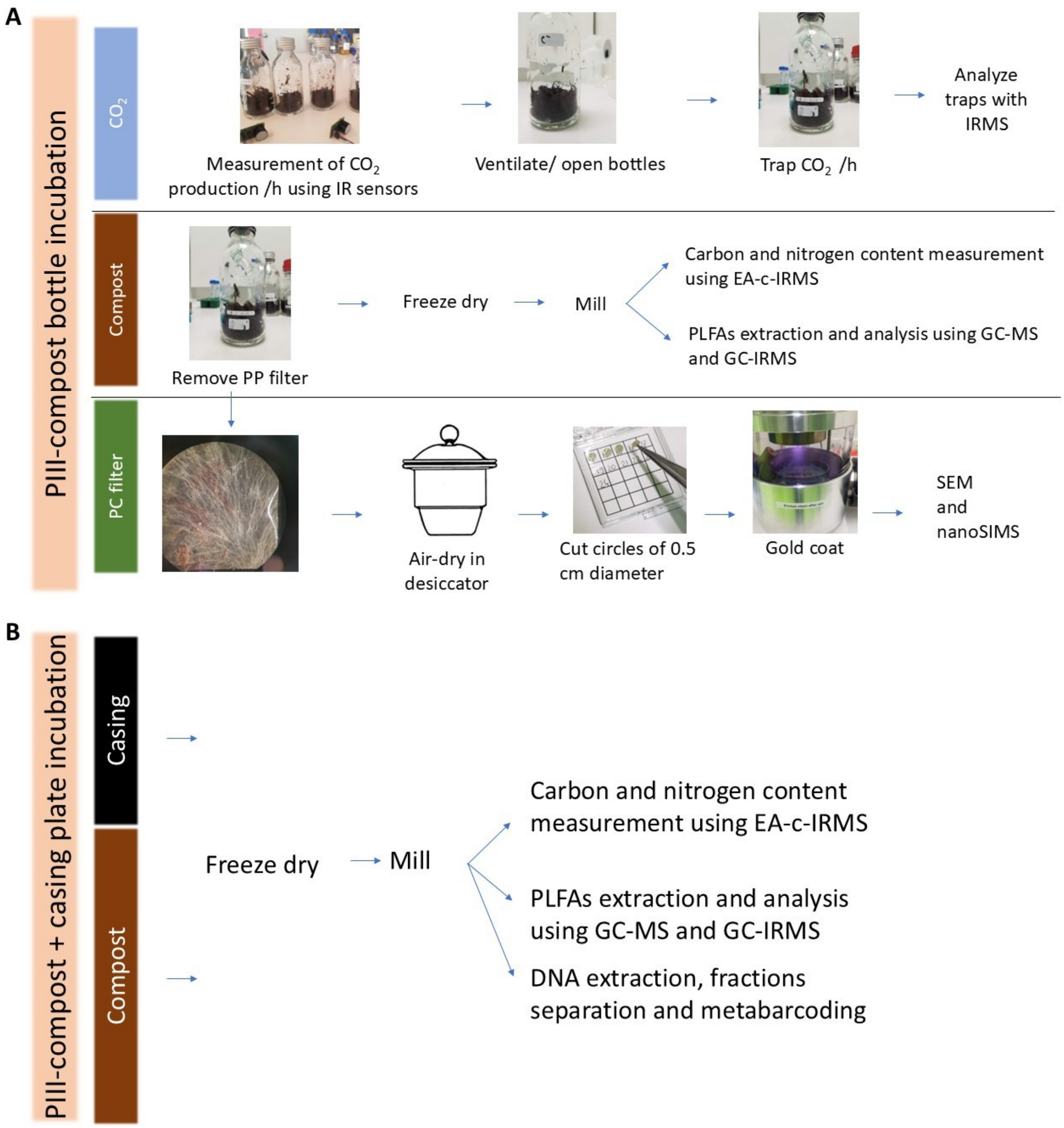


**Figure S3.** Schematic representation of measurements and sample preparation during sampling of compost-based incubations. A: Sampling protocol for the pIII-compost based bottle incubations. The order followed was from top ($CO_2$) to bottom (PP filter). PC = polycarbonate. B: Sampling protocol for pIII-compost and casing-based square plate incubations. For these samples, $CO_2$ and nanoSIMS analysis were not performed, while SIP-PLFA was added as an extra analysis of the substrate (compost and casing).

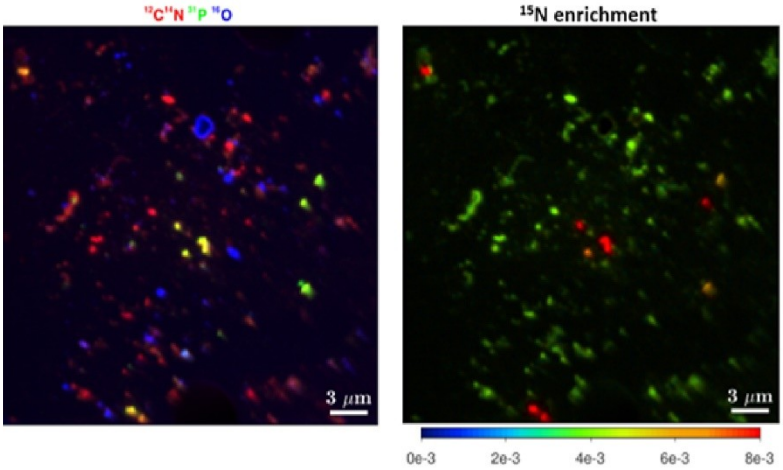


**Figure S4.** Identification of bacteria through nanoSIMS imaging. Left panel: RGB image overlaying the signals from $^{12}C^{14}N$, $^{31}P$, and $^{16}O$, respectively. Bacterial cells were identified according to their size of less than 1.5 µm and the co-localised signals of CN and P (orange and yellow), whereas oxalic acid (blue signal) pieces were excluded from the analysis. Right panel: $^{15}N$ enrichment degree (orange to red hues represent cells that assimilated $^{15}N$). Where possible, this signal was used to confirm the particle identity.

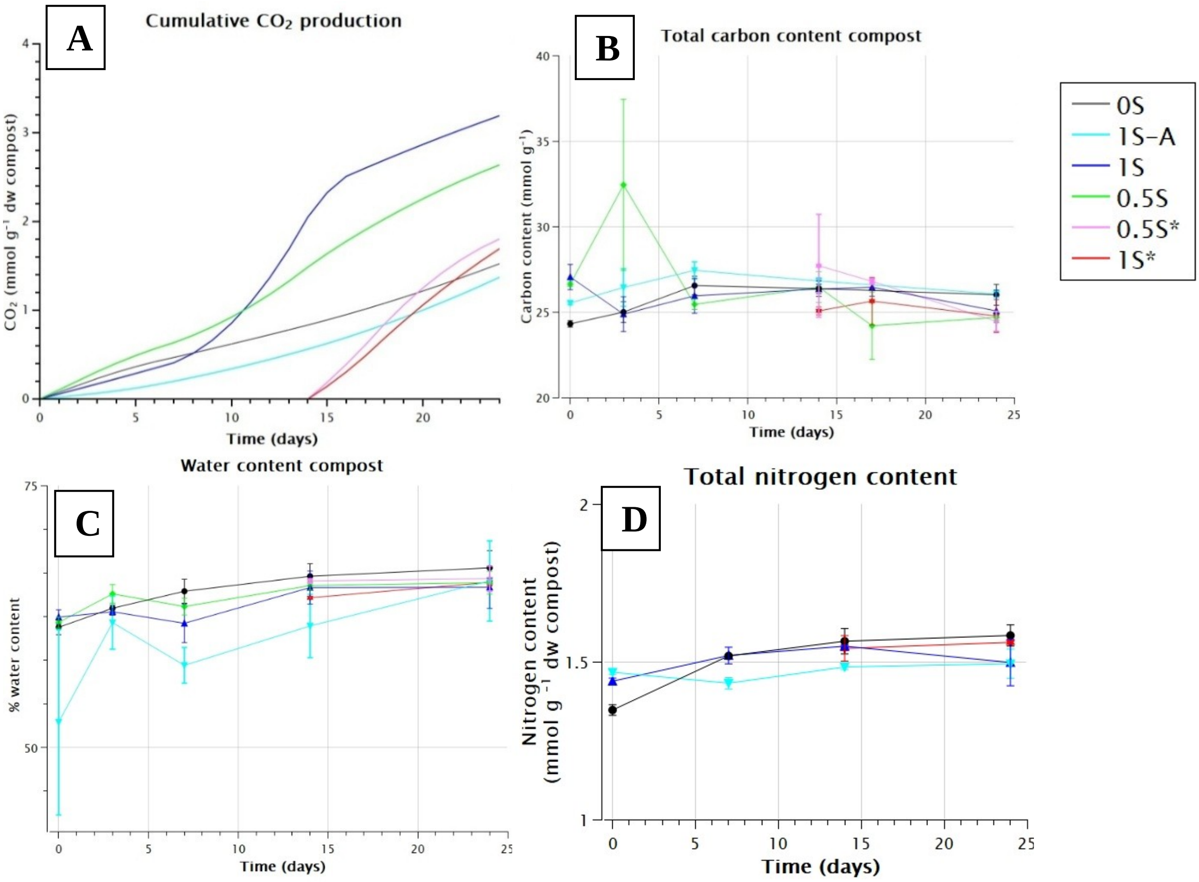


**Figure S5.** Physiochemical measurements of the compost. A: Total $CO_2$ produced during the incubation period, obtained through interpolation of the $CO_2$ production rates at 0, 3, 7, 14, 17, and 24 days. B: Total carbon content of the compost. C: Percentage of water content of the compost (weight %); D: Total nitrogen content of the compost. Error bars represent the minimum and maximum values (n = 2).

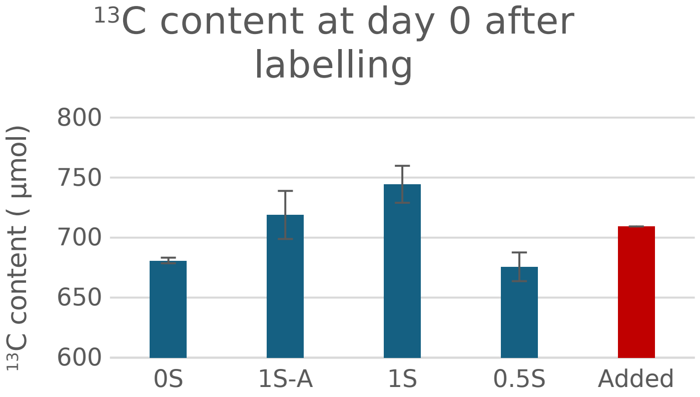


**Figure S6.** Label incorporation test. Amount of $^{13}C$ measured at day 0 under different conditions (blue) against the expected value (red).

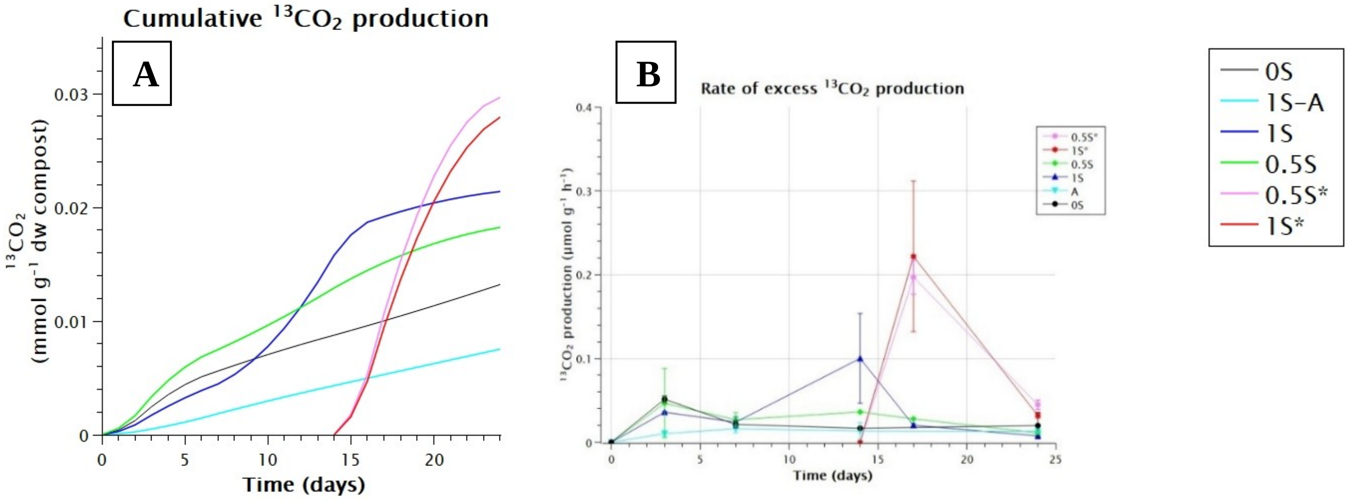


**Figure S7.** $^{13}C$ excess in produced $CO_2$. A: Cumulative $^{13}CO_2$ production over time calculated by interpolating the average rate of glucose respiration for each day from the measured rates. B: Production rate of $^{13}CO_2$ as measured by the $CO_2$ sensor. The $^{13}C$% values were derived from the NaOH traps. Error bars represent standard deviation (SD) for n = 3; where three measurements were not possible, the minimum and maximum values are shown (n = 2).

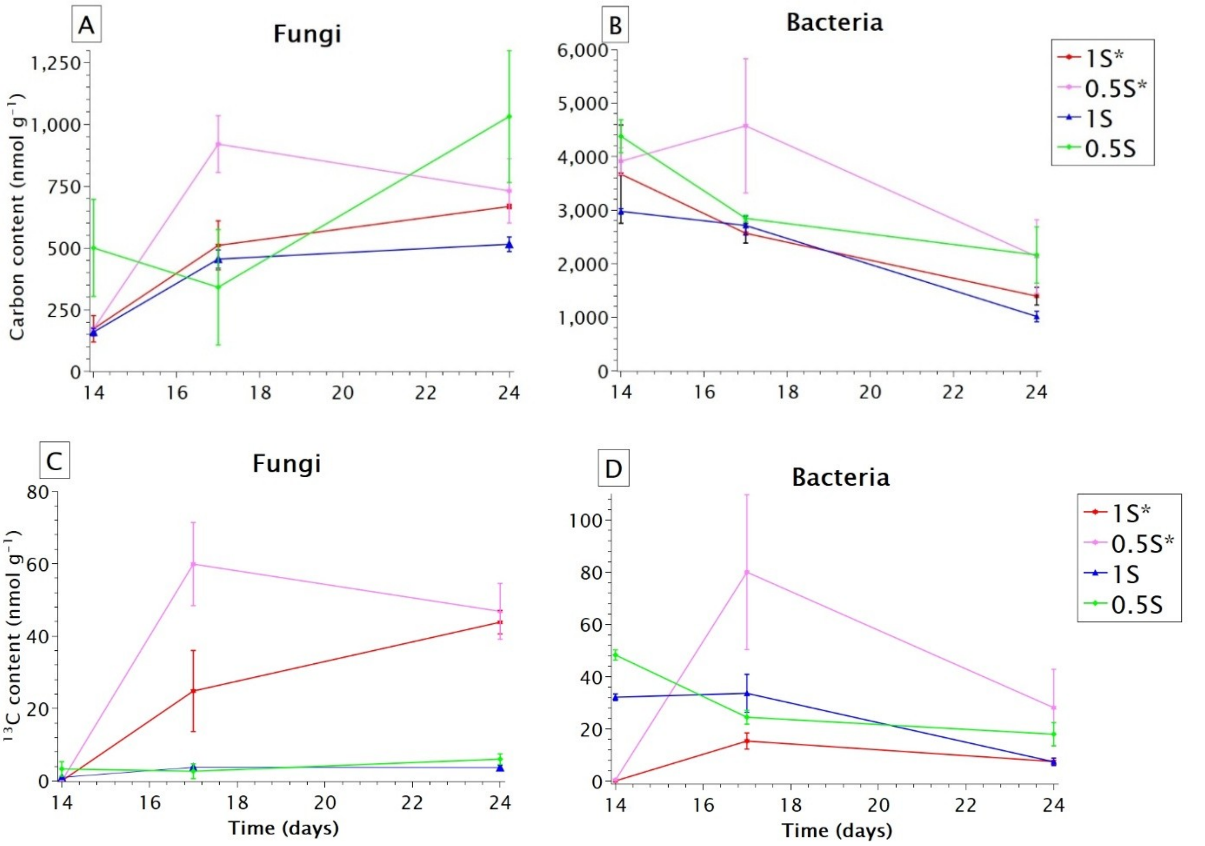


**Figure S8.** PLFA total C concentrations and $^{13}C$ excess for fungal and bacterial biomarkers in parallel experiments. A: Total C in fungal biomarker. B: Total C in bacterial biomarkers. C: $^{13}C$ excess in fungal biomarkers. D: $^{13}C$ excess in bacterial biomarkers. For * conditions, $^{13}C$-glucose was added on day 14. The error bars represent the minimum and maximum values (n = 2).

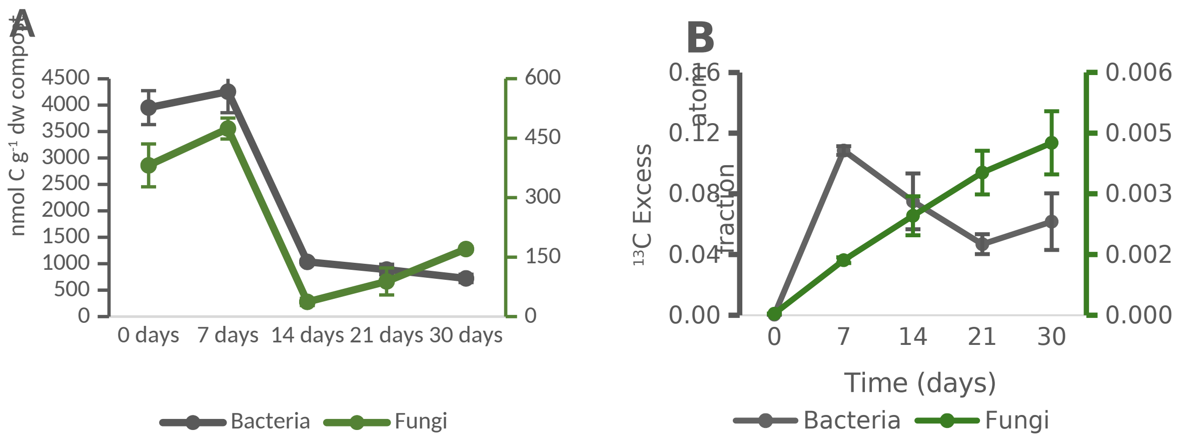

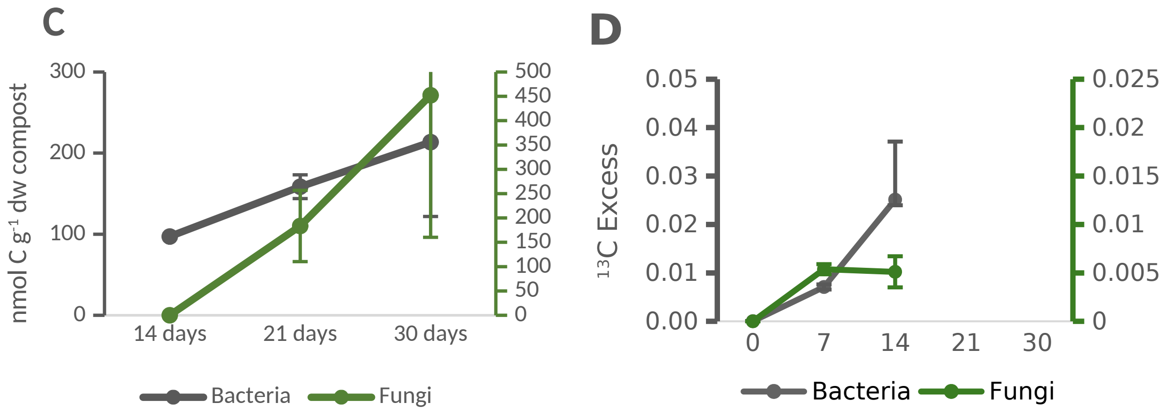


**Figure S9.** Carbon content and tracer excess of bacterial and fungal PLFAs in the compost of the square-dish incubation. A: Concentrations of bacterial and fungal biomarkers in compost. B: Excess $^{13}C$-atom fraction of PLFAs from compost. C: Concentrations of bacterial and fungal biomarkers in the casing layer added on top of phase III compost on day 14. D: Excess $^{13}C$-atom fraction of PLFAs from casing. The error bars indicate the range between the minimum and maximum values (n = 2). The left y-axis shows the $^{13}C$ excess of bacterial PLFAs, while the right y-axis shows the $^{13}C$ excess of fungal PLFAs.

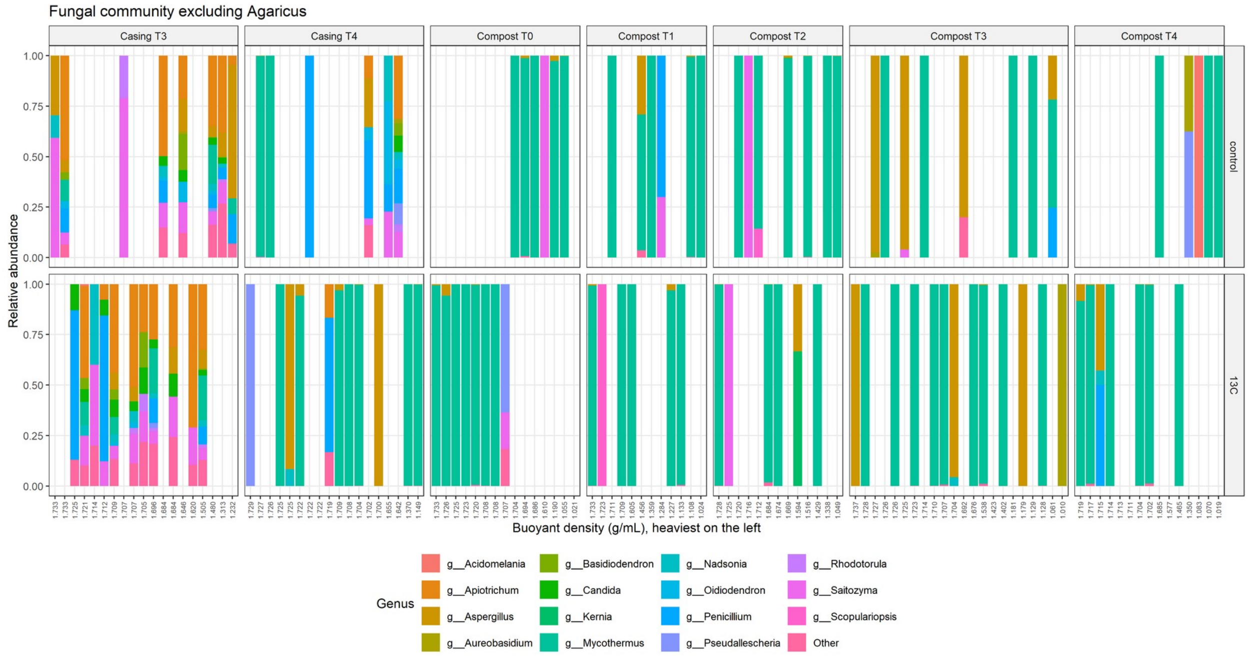


**Figure S10.** Fungal community composition across the CsCl density gradient after excluding *Agaricus*. Panels, axes and density conventions are as in Figure 5, and relative abundances were recalculated after exclusion. *Agaricus* accounts for the large majority of fungal reads, so removing it exposes the remaining community, which is dominated by *Mycothermus thermophilus*. Taxonomy was assigned against a multi-kingdom reference, so plant reads originating from wheat straw are identified as plant and excluded rather than counted as fungal.

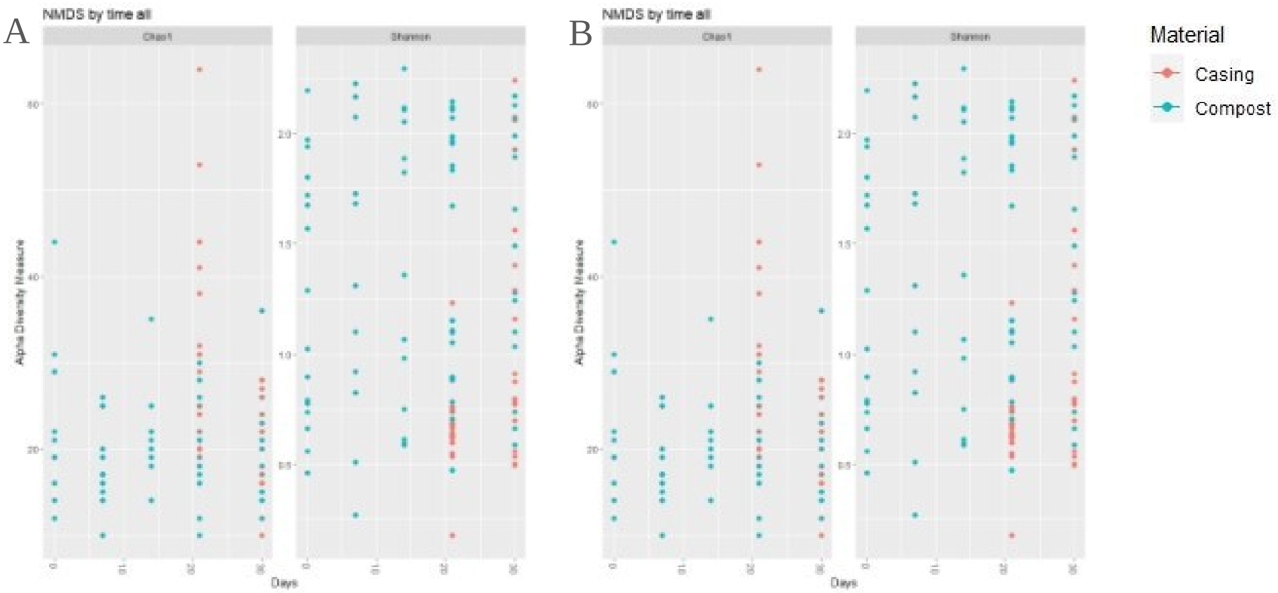


**Figure S11.** Alpha diversity of fungal communities over time in compost and casing substrates. The left panels display species richness estimated using the Chao1 index, while the right panels show the Shannon diversity index, reflecting both the richness and evenness of the microbial communities. A: excludes the genus *Agaricus* from the analysis. B: includes all detected taxa. Data points represent individual samples collected after 0, 10, 20, or 30 days of incubation. The compost substrate generally showed lower alpha diversity than the casing substrate, particularly in the later stages of incubation, with notable differences observed when *Agaricus* was excluded from the analysis.

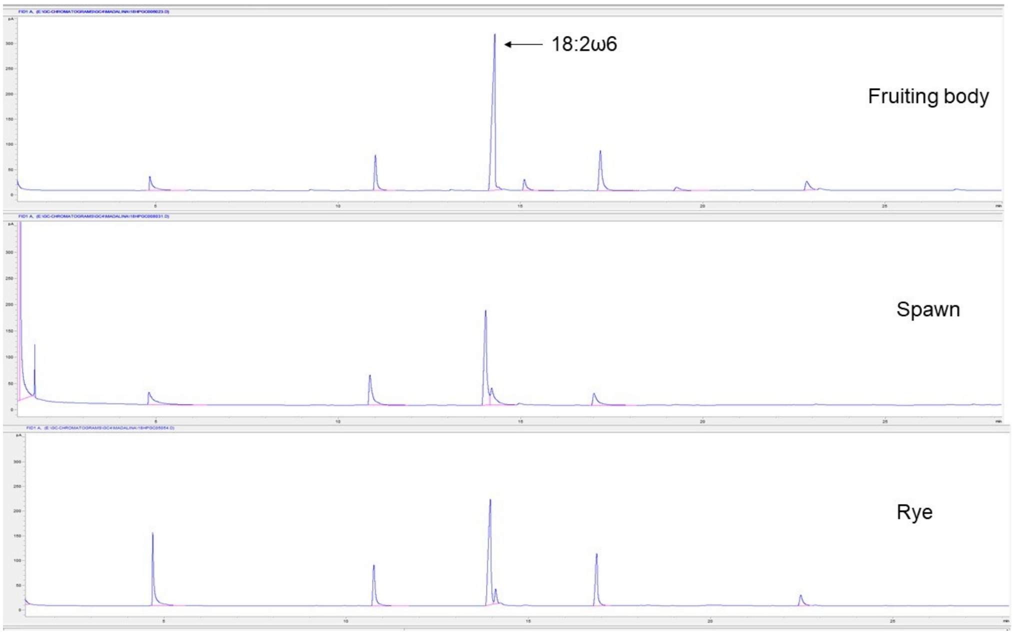


**Figure S12.** Chromatograms of PLFAs extracted from *A. bisporus* fruiting body (Mushroom), spawn, and rye grains. The elution times can differ between runs with ± 1 min.

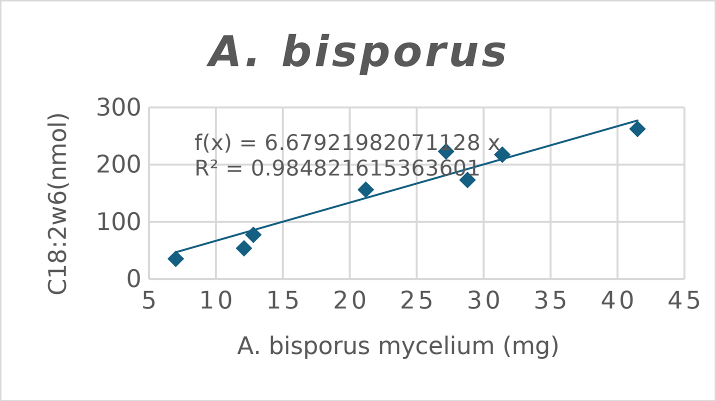


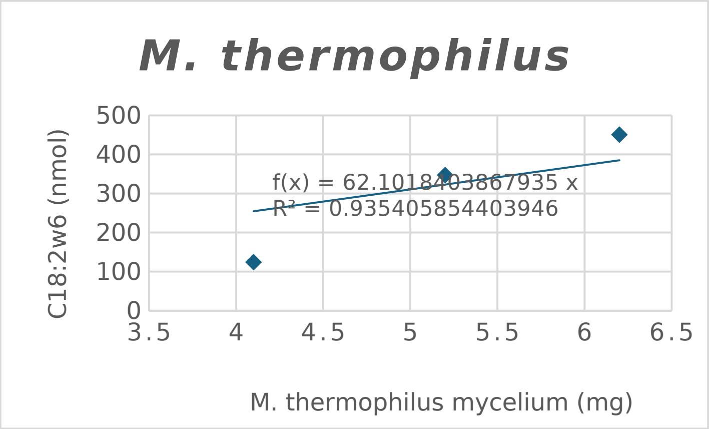


**Figure S13.** Correlation between PLFA content (nmol) and the mass of mycelium used for extraction in the two dominant fungal species in compost: *Agaricus bisporus* (top, n = 8) and *Mycothermus thermophilus* (bottom, n = 3). Lines are least-squares fits constrained through the origin, since a culture of zero biomass contains no PLFA; the slopes are 6.7 and 62.1 nmol C18:2ω6,9c per mg dry weight respectively. One *A. bisporus* culture was excluded because it may have contained residual inoculum. Note that the two species were cultured over non-overlapping biomass ranges, 7.0 to 41.5 mg for *A. bisporus* and 4.1 to 6.2 mg for *M. thermophilus*.

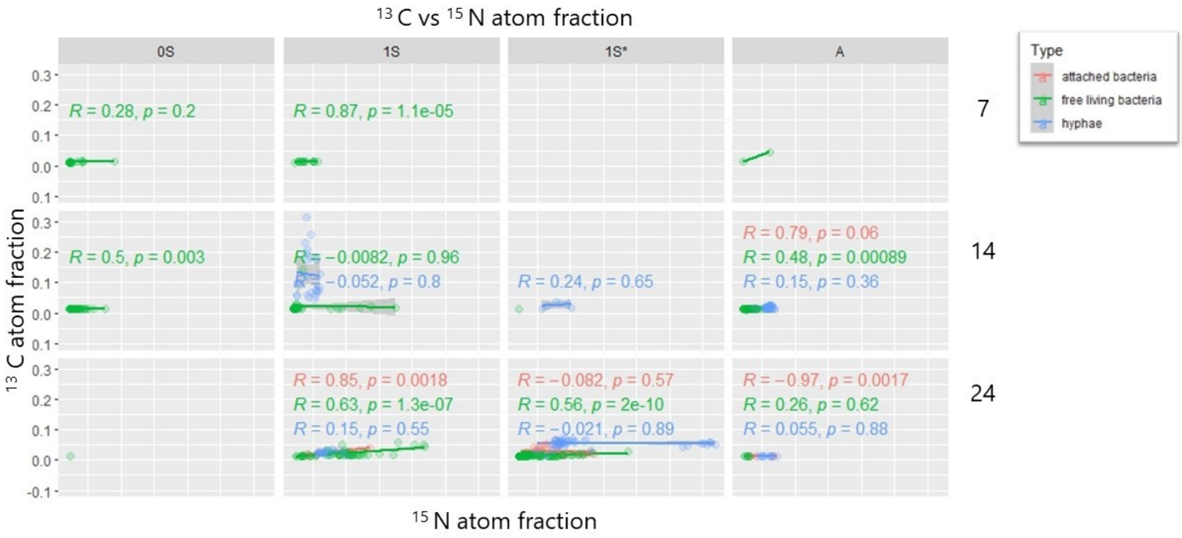


**Figure S14.** nanoSIMS-imaged $^{13}$C and $^{15}$N atom fraction correlation between bacteria and hyphae. Numbers on the left represent sampling times (7, 14 and 24 days of incubation). The top panels indicate the incubation conditions: 0S, no *A. bisporus* spawn; 1S, compost with *A. bisporus*; 1S-A, autoclaved compost with *A. bisporus*; 1S*, same as 1S with the label added after 14 days of incubation.

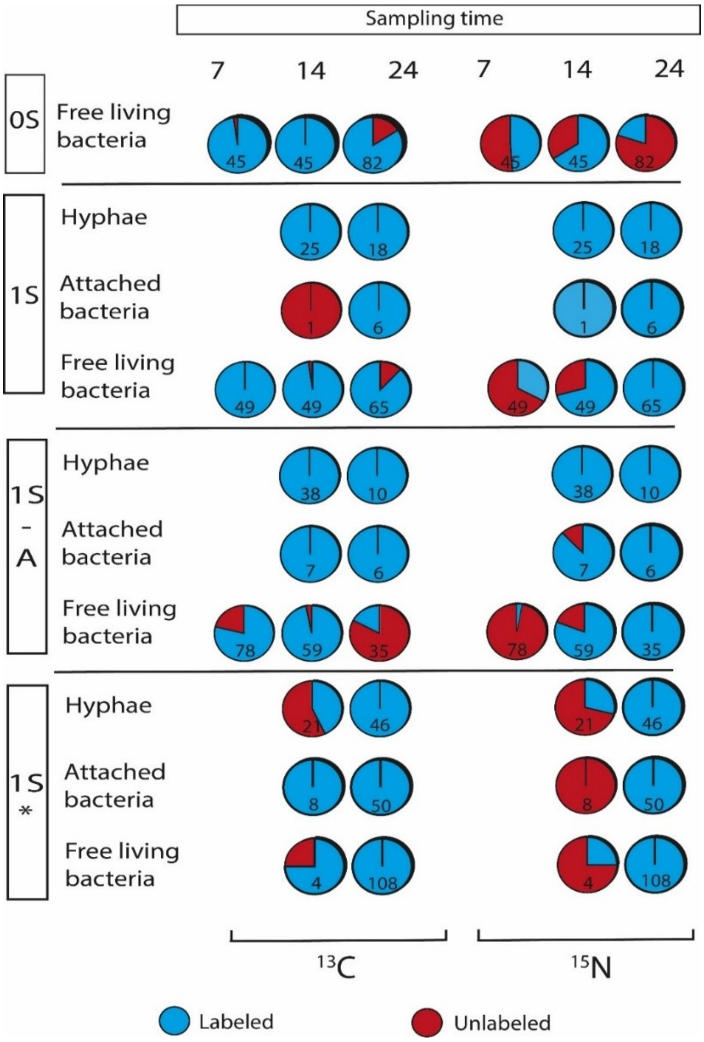


**Figure S15.** NanoSIMS-imaged cell labelling data. The pie charts display the proportion of labelled (blue) and unlabelled (red) cells for both $^{13}C$ and $^{15}N$ isotopes at different sampling times (7, 14, and 24 days) and treatments (0S, 1S, 1S-A, and 1S*). The numbers in each pie chart represent the total number of cells imaged at each respective time point and treatment. The blue segments indicate cells that incorporated the labelled isotopes ($^{13}C$ or $^{15}N$), whereas the red segments represent cells that remained unlabelled. The breakdown is shown separately for free-living bacteria, hyphae, and bacteria attached to hyphae. The treatments varied according to the conditions applied: 0S served as the control without *Agaricus bisporus*, 1S represented the presence of *A. bisporus*, 1S-A was *A. bisporus* grown in autoclaved compost, and 1S* involved *A. bisporus* with an isotope label added after 14 days of incubation.

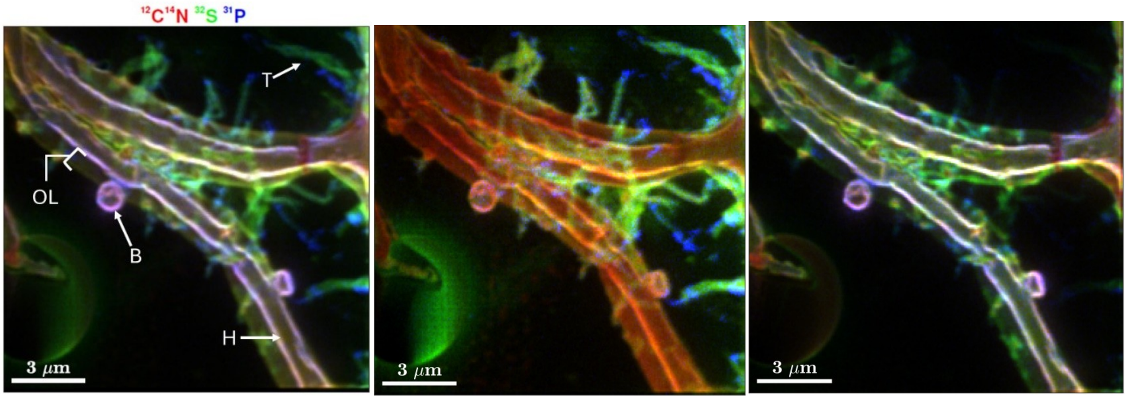


**Figure S16**. NanoSIMS imaging of fungal–bacterial co-occurrence spots. Red, green, and blue (RGB) image of organic carbon and nitrogen (red), sulphur (green), and phosphorus (blue). The same field of view (FOV) can be observed at different depths. Left: cumulative image of all 400 planes measured; middle: first 50 planes; right: plane 200 to 400. Legend: B = bacterial cell; H = hyphal filament; T = sulphur-rich threads; OL = outer layer of the hyphae.

### Supplementary Results: density separation

Density separation was attempted so that $^{13}C$-labelled DNA could be assigned to individual taxa. The labelling achieved was too low for this to be feasible. The calculation below is likely to apply to stable-isotope probing in compost more generally.

The buoyant density of fully $^{13}C$-labelled DNA lies approximately 0.036 g/mL above that of unlabelled DNA, and the shift scales with the proportion of carbon replaced. In this incubation the fungal biomarker reached 0.52 atom% excess $^{13}C$ (Fig. S9B, D), corresponding to a shift of about 0.0002 g/mL. Adjacent fractions within a gradient were separated by a median of 0.0041 g/mL, so moving fungal DNA by even one fraction step would have required approximately 11 atom% excess, and a conventional 0.03 g/mL separation approximately 83 atom%. The cause is dilution: the glucose added represented a small fraction of the carbon already present in the compost, so the fungi drew most of their carbon from the substrate rather than from the tracer.

PERMANOVA on Bray–Curtis distances across the 64 in-gradient fractions returned substrate as the only significant term ($R^2$ = 0.070, F = 4.79, p = 0.034), with neither treatment (p = 0.225) nor buoyant density (p = 0.481) significant (Fig. S17).

This is a limitation of detection rather than evidence that the fungi were inactive. PLFA-SIP on the same incubations shows the excess $^{13}C$ atom fraction of fungal biomarkers rising throughout the experiment in both compost and casing (Fig. S9B, D).

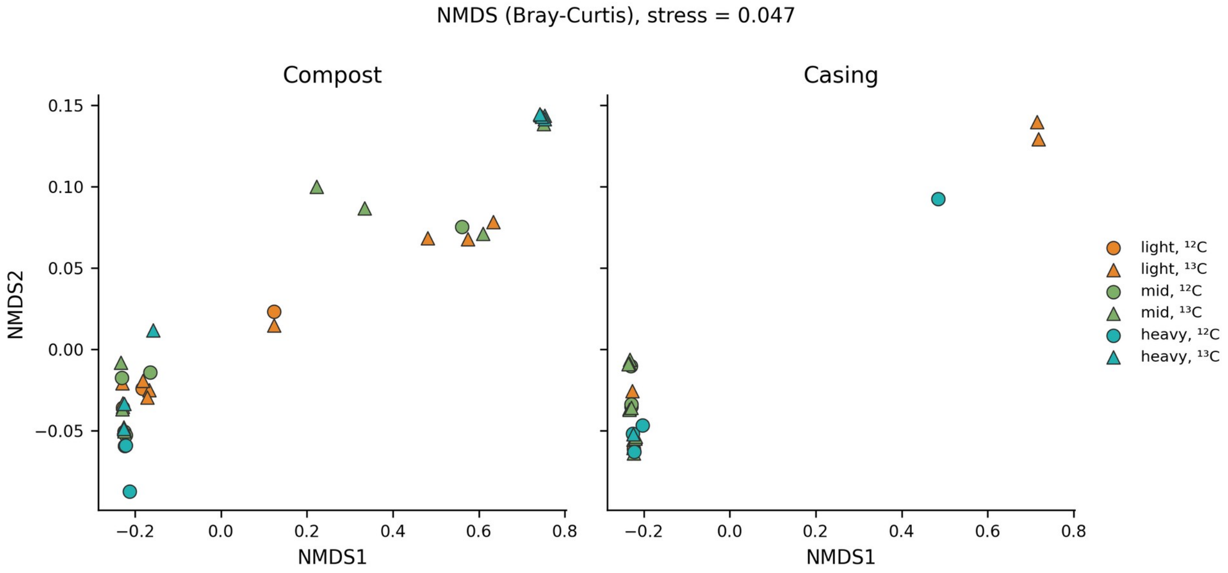


**Figure S17.** Non-metric multidimensional scaling (NMDS) of fungal community composition, Bray–Curtis distances, for the 64 sequenced fractions falling inside the CsCl gradient. Symbols distinguish unlabelled (circles) from $^{13}C$-labelled (triangles) gradients and colour indicates density class. Compost and casing are shown separately. Samples do not separate by treatment or by density; PERMANOVA on the same distances returns substrate as the only significant term ($R^2 = 0.070$, $p = 0.034$), with treatment ($p = 0.225$) and buoyant density ($p = 0.481$) both non-significant.

### Supplementary Tables S1–S5

**Table S1.** PLFA biomarkers used for identification and biomass calculation of microbial communities.

| *Organism* | PLFA | Reference |
|---|---|---|
| *Gram-positive bacteria* | i14:0, i15:0, ai15:0, i17:0, ai17:0 | (Zelles 1997, 1999) |
| *Gram-negative bacteria* | cy17:0, cy19:0, C16:1w7 | (Zelles 1997, 1999) |
| *Bacteria (general)* | 18:1ω7c | (Zelles 1997, 1999) |
| *Actinobacteria* | 10Me-C16:0 | (Zelles 1999) |
| *Fungi* | 18:2ω6,9c | (Matcham et al. 1985; Frostegard and Baath 1996) |

#### Statistical analysis tables

Each row gives the omnibus test (one-way ANOVA F, or Kruskal–Wallis $\chi^2$ where variances were heterogeneous; t = two-sample t-test for the re-fed 1S* comparisons) and its p-value; the final column lists only the comparisons that were significant after Bonferroni adjustment ($p < 0.05$). Non-significant pairwise comparisons (all $p \approx 1$) are omitted. Significant omnibus p-values are shaded green. Treatments: A = autoclaved; 0S = no spawn; 0.5S = half spawn density; 1S = standard spawn density; * = isotope/glucose re-applied at day 14. Time points T0–T5 = days 0, 3, 7, 14, 17, 24.

**Table S2.** $CO_2$ and bulk measurements (excess $^{13}C$, total carbon, total $CO_2$)

| $CO_2$ — Excess $^{13}C$ (mmol), per time point |
|---|

| | | | |
|---|---|---|---|
| **Day** | **F / χ²** | **p** | **Significant pairwise comparisons (Bonferroni p)** |
| 0 | — | — | — |
| 3 | 23.49 | **0.005** | A–0.5S 0.008; A–0S 0.030 |
| 7 | 1.674 | 0.249 | none |
| 14 | 13.06 (χ²) | **0.023 (KW)** | 0.5S*–1S 0.034 |
| 17 | 4.383 | 0.073 | none |
| 24 | 5.779 | **0.015** | 0.5S–0.5S* 0.041; 0.5S*–1S 0.038 |

| **$CO_2$ — Total $CO_2$ produced (mmol), per time point** | | | |
|---|---|---|---|
| **Day** | **F / χ²** | **p** | **Significant pairwise comparisons (Bonferroni p)** |
| 0 | — | — | — |
| 3 | 18.03 | **0.009** | A–0.5S 0.012 |
| 7 | 1.945 | 0.201 | none |
| 14 | 11.07 (χ²) | 0.05 (KW) | none |
| 17 | 1.53 | 0.315 | none |
| 24 | 0.8 | 0.558 | none |

| **Bulk compost — Excess $^{13}C$ (mmol), per time point** | | | |
|---|---|---|---|
| **Day** | **F / χ²** | **p** | **Significant pairwise comparisons (Bonferroni p)** |
| 0 | 2.875 | 0.167 | none |
| 3 | 8.123 | **0.035** | none significant after correction |
| 7 | 0.396 | 0.763 | none |
| 14 | 18.29 | **0.001** | A–0.5S* 0.012; A–1S* 0.014; 0.5S–0.5S* 0.016; 0.5S–1S* 0.020; 0.5S*–0S 0.035; 0.5S*–1S 0.010; 0S–1S* 0.044; 1S–1S* 0.012 |
| 17 | 46.69 | **0.001** | 0.5S–0.5S* 0.012; 0.5S–1S* 0.004; 0.5S*–1S 0.016; 1S–1S* 0.005 |
| 24 | 9.839 | **0.007** | A–1S* 0.015 |

| Bulk compost — Total carbon (mmol), per time point | | | |
|---|---|---|---|
| **Day** | **F / χ²** | **p** | **Significant pairwise comparisons (Bonferroni p)** |
| 0 | 3.348 | 0.137 | none |
| 3 | 6.167 (χ²) | 0.104 (KW) | none |
| 7 | 16.67 | **0.010** | A–0.5S 0.036; A–0S 0.020; A–1S 0.029 |
| 14 | 2.588 | 0.139 | none |
| 17 | 1.577 | 0.327 | none |
| 24 | 0.952 | 0.511 | none |

**Table S3.** Nitrogen measurements (excess $^{15}N$ and total nitrogen)

Values transcribed from the existing table (source spreadsheet not yet located). Treatment 1S* (days 14 and 24 only) compared by two-sample t-test.

| Excess $^{15}N$ (mmol) — per time point (between conditions) | | | |
|---|---|---|---|
| **Day** | **F-value** | **p** | **Significant pairwise comparisons (Bonferroni p)** |
| 0 | 0.467 | 0.666 | none |
| 7 | 59.036 | **0.004** | A–0S 0.0076; 1S–A 0.0077 |
| 14 | 48.060 | **0.001** | 1S*–0S 0.0031; 1S*–1S 0.0027; 1S*–A 0.0163 |
| 24 | 9.127 | **0.029** | A–0S 0.0440 |

| Excess $^{15}N$ (mmol) — per condition (between time points) | | | |
|---|---|---|---|
| **Cond.** | **F / T** | **p** | **Significant time-pairs (Bonferroni p)** |
| 1S | 129.27 | **0.0002** | T0–T2 0.0004; T0–T3 0.0005; T0–T5 0.0008 |
| 1S* | -6.638 (t) | **0.022 (t)** | T3–T5 0.022 |
| 0S | 403.1 | **<0.001** | T0–T2 <0.001; T0–T3 0.0001; T0–T5 0.0001 |
| A | 28.11 | **0.004** | T0–T3 0.0071; T0–T5 0.0105 |

| Total nitrogen (mmol) — per time point | | | |
|---|---|---|---|
| **Day** | **F-value** | **p** | **Significant pairwise comparisons (Bonferroni p)** |

| | | | |
|---|---|---|---|
| 0 | 6.804 | 0.077 | none |
| 7 | 1.530 | 0.348 | none |
| 14 | 1.607 | 0.321 | none |
| 24 | 0.312 | 0.817 | none |

| **Total nitrogen (mmol) — per condition** | | | |
|---|---|---|---|
| **Cond.** | **F / T** | **p** | **Significant time-pairs (Bonferroni p)** |
| 1S | 0.976 | 0.487 | none |
| 1S* | -2.117 (t) | 0.169 (t) | none |
| 0S | 1.157 | 0.429 | none |
| A | 3.777 | 0.116 | none |

**Table S4.** Fungal PLFA results (excess $^{13}C$ and total carbon)

| **Fungal PLFA — Excess $^{13}C$ — per condition (between time points)** | | | |
|---|---|---|---|
| **Cond.** | **F-value** | **p** | **Significant time-pairs (Bonferroni p)** |
| 1S | 35.19 | **0.0002** | T0–T1 0.002; T0–T2 0.001; T1–T3 0.003; T2–T3 0.001; T1–T4 0.019; T2–T4 0.006; T1–T5 0.018; T2–T5 0.006 |
| 0.5S | 3.695 | 0.071 | none |
| 0S | 457.1 | **<0.001** | T0–T1 <0.001; T0–T2 <0.001; T0–T3 <0.001; T1–T3 0.011; T2–T3 0.014; T0–T5 0.0001; T1–T5 0.0002; T2–T5 0.0002; T3–T5 0.002 |
| A | 2.874 | 0.139 | none |
| 1S* | 10.67 | **0.043** | none significant after correction |
| 0.5S* | 15.55 | **0.026** | T3–T4 0.039 |

| **Fungal PLFA — Excess $^{13}C$ — per time point (between conditions)** | | | |
|---|---|---|---|
| **Day** | **F-value** | **p** | **Significant comparisons (Bonferroni p; t = t-test)** |
| 0 | 34.29 | **0.009** | 0.5S–0S 0.013; 0S–1S 0.023 |
| 3 | 8.636 | 0.057 | 1S–A 0.010 (t) |

| | | | |
|---|---|---|---|
| 7 | 17.51 | **0.022** | 0.5S–1S 0.036; 1S–A 0.022 (t) |
| 14 | 12.04 | **0.037** | 1S–1S* 0.018 (t) |
| 17 | — | — | none |
| 24 | 2.238 | 0.254 | 1S–1S* 0.007 (t); 0.5S–0.5S* 0.035 (t) |

| **Fungal PLFA — Total carbon (nmol g⁻¹) — per condition (between time points)** | | | |
|---|---|---|---|
| **Cond.** | **F-value** | **p** | **Significant time-pairs (Bonferroni p)** |
| 1S | 21.31 | **0.0009** | T0–T2 0.039; T1–T3 0.005; T2–T3 0.001; T2–T4 0.015; T2–T5 0.030 |
| 0.5S | 3.065 | 0.103 | none |
| 0S | 2.311 | 0.191 | none |
| A | 4.250 | 0.072 | none |
| 1S* | 14.98 | **0.027** | T3–T5 0.038 |
| 0.5S* | 161.6 | **0.001** | T3–T4 0.002; T3–T5 0.002 |

| **Fungal PLFA — Total carbon (nmol g⁻¹) — per time point (between conditions)** | | | |
|---|---|---|---|
| **Day** | **F-value** | **p** | **Significant comparisons (Bonferroni p; t = t-test)** |
| 0 | 17.58 | **0.022** | 0.5S–1S 0.033 |
| 3 | 4.604 | 0.122 | 1S–A 0.037 (t) |
| 7 | 13.70 | **0.031** | 0.5S–1S 0.041; 1S–A 0.034 (t) |
| 14 | 4.63 | 0.121 | none |
| 17 | — | — | none |
| 24 | 24.20 | **0.014** | 0.5S–0S 0.019; 1S–1S* 0.035 (t) |

**Table S5.** Bacterial PLFA results (excess $^{13}C$ and total carbon)

| **Bacterial PLFA — Excess $^{13}C$ — per condition (between time points)** | | | |
|---|---|---|---|
| **Cond.** | **F-value** | **p** | **Significant time-pairs (Bonferroni p)** |
| 1S | 110.49 | **<0.001** | T0–T1 <0.001; T0–T2 <0.001; T1–T3 <0.001; T2–T3 <0.001; T1–T4 <0.001; T2–T4 <0.001; T1–T5 <0.001; T2–T5 <0.001 |

| | | | |
|---|---|---|---|
| 0.5S | 101.4 | **<0.001** | T0–T1 <0.001; T0–T2 0.001; T1–T2 0.001; T0–T3 0.010; T1–T3 <0.001; T1–T4 <0.001; T2–T4 0.008; T1–T5 <0.001; T2–T5 0.004 |
| 0S | 80.63 | **<0.001** | T0–T1 <0.001; T0–T2 <0.001; T0–T3 0.002; T2–T3 0.007; T0–T5 0.003; T2–T5 0.005 |
| A | 20.31 | **0.003** | T0–T2 0.005; T1–T2 0.008; T2–T3 0.034; T2–T5 0.011 |
| 1S* | 15.63 | **0.026** | T3–T4 0.017 |
| 0.5S* | 5.960 | 0.090 | none |

| **Bacterial PLFA — Excess $^{13}C$ — per time point (between conditions)** | | | |
|---|---|---|---|
| **Day** | **F-value** | **p** | **Significant comparisons (Bonferroni p; t = t-test)** |
| 0 | 3.598 | 0.160 | none |
| 3 | 9.046 | 0.054 | 1S–A 0.007 (t) |
| 7 | 60.72 | **0.004** | 0.5S–0S 0.009; 0.5S–1S 0.006; 1S–A 0.021 (t) |
| 14 | 29.04 | **0.011** | 0.5S–0S 0.028; 0S–1S 0.017; 1S–1S* 0.002 (t); 0.5S–0.5S* 0.002 (t) |
| 17 | — | — | none |
| 24 | 460.5 | **<0.001** | 0.5S–0S <0.001; 0S–1S <0.001 |

| **Bacterial PLFA — Total carbon (nmol $g^{-1}$) — per condition (between time points)** | | | |
|---|---|---|---|
| **Cond.** | **F-value** | **p** | **Significant time-pairs (Bonferroni p)** |
| 1S | 51.27 | **0.0001** | T0–T1 0.015; T0–T2 0.001; T1–T3 0.007; T2–T3 0.001; T1–T4 0.006; T2–T4 0.001; T1–T5 0.001; T2–T5 <0.001 |
| 0.5S | 19.93 | **0.001** | T0–T2 0.023; T0–T4 0.007; T1–T4 0.017; T0–T5 0.003; T1–T5 0.008 |
| 0S | 10.59 | **0.012** | T0–T2 0.018 |
| A | 153.4 | **<0.001** | T0–T2 0.007; T1–T2 0.008; T0–T3 0.001; T1–T3 0.001; T2–T3 <0.001; T0–T5 <0.001; T1–T5 <0.001; T2–T5 <0.001 |
| 1S* | 3.04 | 0.190 | none |
| 0.5S* | 3.27 | 0.177 | none |

| Bacterial PLFA — Total carbon (nmol $g^{-1}$) — per time point (between conditions) | | | |
|---|---|---|---|
| **Day** | **F-value** | **p** | **Significant comparisons (Bonferroni p; t = t-test)** |
| 0 | 19.74 | **0.019** | 0.5S–1S 0.040; 0S–1S 0.034 |
| 3 | 3.999 | 0.143 | 1S–A 0.036 (t) |
| 7 | 49.83 | **0.005** | 0.5S–0S 0.018; 0.5S–1S 0.007; 1S–A 0.006 (t) |
| 14 | 83.85 | **0.002** | 0.5S–0S 0.010; 0S–1S 0.003; 1S–A 0.006 (t) |
| 17 | — | — | none |
| 24 | 157.5 | **0.001** | 0.5S–0S 0.002; 0S–1S 0.002 |
